# Machine learning identifies routine blood tests as accurate predictive measures of pollution-dependent poor cognitive function

**DOI:** 10.1101/2025.01.10.632396

**Authors:** Hamish Johnson, James Longden, Gary Cameron, Gordon D. Waiter, Fergal M. Waldron, Jenna M. Gregory, Holly Spence

## Abstract

**Background:** Several modifiable risk factors for dementia and related neurodegenerative diseases have been identified including education level, socio-economic status, and environmental exposures – however, how these population-level risks relate to individual risk remains elusive. To address this, we assess over 450 potential risk factors in one deeply clinically and demographically phenotyped cohort using random forest classifiers to determine predictive markers of poor cognitive function. This study aims to understand early risk factors for dementia by identifying predictors of poor cognitive performance amongst a comprehensive battery of imaging, blood, atmospheric pollutant and socio-economic measures.

**Methods:** Random forest modelling was used to determine significant predictors of poor cognitive performance in a cohort of 324 individuals (age 61.6 ± 4.8 years; 150 males, 174 females) without extant neurological disease. 457 features were assessed including brain imaging measures of volume and iron deposition, blood measures of anaemia, inflammation, and heavy metal levels, social deprivation indicators and atmospheric pollution exposure.

**Results:** Routinely assessed markers of anaemia including mean corpuscular haemoglobin concentration were identified as robust predictors of poor general cognition, where both extremes (low and high) were associated with poor cognitive performance. The strongest, most consistent predictors of poor cognitive performance were environmental measures of atmospheric pollution, in particular, lead, carbon monoxide, and particulate matter. Feature analysis demonstrated a significant negative relationship between low mean corpuscular haemoglobin concentration and high levels of atmospheric pollutants highlighting the potential of routinely assessed blood tests as a predictive measure of pollution-dependent cognitive functioning, at an individual level.

**Conclusions:** Taken together, these data demonstrate how routine, inexpensive medical testing and local authority initiatives could help to identify and protect at-risk individuals. These findings highlight the potential to identify individuals for targeted, cost effective medical and social interventions to improve population cognitive health.

**Graphical Abstract:** 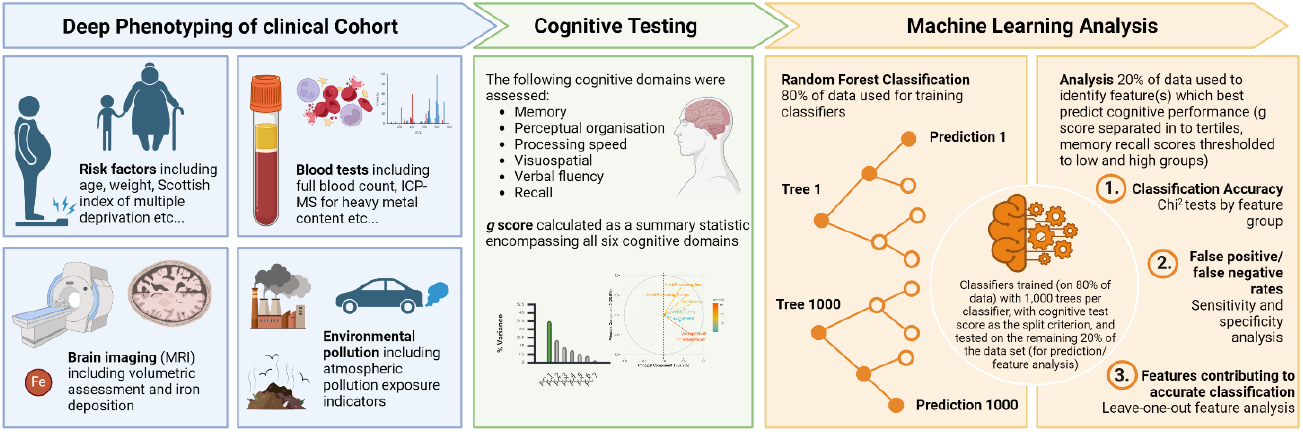

## Introduction

Many risk factors for dementia have been identified including hypertension and diabetes, however there is often conflicting results across studies (Jia et al., 2020; Dintica & Yaffe 2022; Livingston et al., 2023; Zhang et al., 2023). The Lancet commission report on dementia prevention highlighted twelve modifiable risk factors which are believed to account for 40% of dementias (Livingston et al., 2023). These included lower education, obesity, low social contact, and air pollution. By looking at modifiable factors that affect incident cognitive performance, we aim to define readily identifiable environmental and health factors responsible for poor cognitive performance to influence policy and improve cognitive health for those most at risk. To do this we assess a comprehensive battery of socio-demographic risk factors, brain imaging markers, blood markers and atmospheric pollutants in a deeply phenotyped cohort of older individuals.

Higher levels of social deprivation have previously been associated with many diseases including an increased risk of dementia (Sakaniwa et al., 2024, Russ et al., 2013). In these studies, characterisation of socio-economic status varies widely from leaving education at an earlier age to social class, and there remains a lack of consensus regarding which specific socio-economic stressors may impact cognitive health most substantially. The current cohort allows the unique opportunity to combine comprehensive socio-demographic information in the form of the Scottish Index for Multiple Deprivation (SIMD; www.SIMD.scot) with other health indicators to identify areas where social change may benefit cognitive health. Whilst identification of specific social deprivation predictors of cognitive health is essential to determining policy and lifestyle change, it is important to also consider imaging or blood predictors that could be used as biomarkers of cognitive health. Brain imaging measures of regional volume and regional iron, as well as blood markers of anaemia and inflammation have previously been shown to have an association with dementia and so are also included in the current study (Kravitz et al., 2010; Tachibana et al., 2024; Hong et al., 2013; Wolters et al., 2019; Spence et al., 2024; Schuff et al., 2009).

Exposure to air pollution has been associated with cognitive health variability with longitudinal studies showing that regardless of region of recruitment (i.e. area of dwelling), levels of particulate matter (PM2.5), carbon monoxide and nitrogen dioxide were associated with cognitive decline and dementia incidence (Peters et al., 2019). Whilst several studies show a positive association between pollutant levels and rates of dementia, some studies showed a lack of association (Tonne et al., 2014) or a negative association (Carey et al., 2018). This highlights the need for further study in a large deeply phenotyped cohort to elucidate the nuanced relationships between pollution and cognitive health.

Using a non-biased approach, here we assess a deeply phenotyped cohort of older individuals with no reported neurological disease to determine the strongest predictors of poor cognitive function to detect those at increased risk of dementia and other neurodegenerative disorders. Phenotyping of this cohort with brain imaging, blood sample testing, cognitive testing and spatially linked Scottish index of multiple deprivation and atmospheric pollution data allows for comprehensive modelling of cognitive predictors. Robust cognitive assessment comprised seven measures over five cognitive domains including memory and recall (immediate and delayed tests), perceptual organisation, processing speed, matrix reasoning (time and score) and verbal fluency. These seven measures were combined via principal component analysis to generate a *g* factor for general cognition (*g*).

(see Van der Maas et al., 2017; Spence et al., 2022). Using this approach, we can gain insights into predictors of general cognitive health. We also analyse these cognitive domains separately where each domain represents a risk factor for dementia subtypes, for example, poorer memory score represents a higher risk for Alzheimer’s type dementia. We establish a random forest modelling approach to allow for non-biased feature identification, determining predictors of poor cognitive function. This study aims to establish risk factors for which routine clinical and local authority intervention could be taken to identify and protect ‘at risk’ populations (Figure 1).

**Figure 1.**
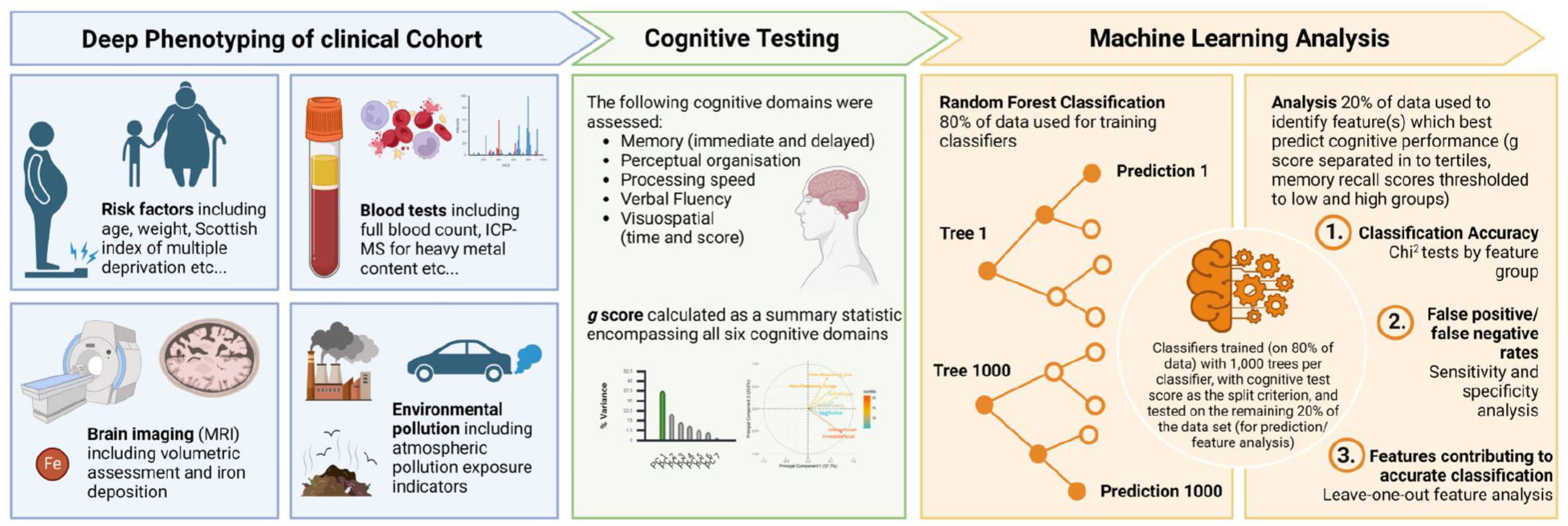
Cohort profiling and random forest modelling workflow

## Methods

### Cohort profile

Participants of the Aberdeen Children of the 1950’s (ACONF) study and their first-generation relatives who had given consent to be recontacted and still lived in Scotland with a community health index (CHI) number were recruited as part of the STratifying Resilience and Depression Longitudinally (STRADL) study (Habota et al., 2019). Ethical approval was previously obtained from the Scotland A Research Ethics Committee (REC reference number 14/55/0039) and the local Research and Development offices. Participants were excluded from this study if they had a self-reported diagnosis of a neurological condition or neurodegenerative disease, or if they did not have MRI data suitable for quantitative susceptibility mapping (QSM). A total of 324 individuals (61.6 ± 4.8 y/o; 150 males, 174 females) were included in this study and measurements were acquired between 2015 and 2017. A series of cognitive tests were performed on day of MRI and blood sampling. These included the Mill Hill Vocabulary test, the controlled word association task, the logical memory subset (immediate and delayed recall) and digit symbol coding task of the Wechsler adult intelligence scale III and the matrix reasoning test of the COGNITO psychometric examination (Habota et al., 2019). Scottish Index of Multiple Deprivation (SIMD) was used as a relative measure of deprivation and was collected from the freely available database (www.simd.scot) based on the known postcode of the participants on the day of cognitive assessment, imaging and blood sampling. The 2016 SIMD measures were used as this was the closest available timepoint to MRI, blood and cognitive testing.

### Imaging

Imaging protocols are described fully by Habota et al., 2019. Briefly, images were acquired on a 3T Philips Achieva TX-series MRI system (Philips Healthcare, Best, Netherlands) with a 32-channel phased-array head coil. 3D T1 weighted fast gradient echo images were acquired with the following parameters: 160 sagittal slices, TR = 8.2 ms, TE = 3.8 ms, TI = 1031 ms, FA = 8°, FOV = 240 mm, matrix size = 240 × 240, voxel size = 1.0 × 1.0 × 1.0 mm^3^, acquisition time = 5 min 38 s. MEGRE images were acquired with the following parameters: 130 axial slices, TR = 31, TE = 7.2/13.4/19.6/25.8 ms, FA = 17°, FOV = 230 mm, matrix size = 384 × 316, Voxel size = 0.3 × 0.3 × 1 mm^3^, acquisition time = 4min 29s. Brain masks were generated using the brain extraction tool function of fsl software (Smith, 2002). Phase images from MEGRE data were processed through a QSM pipeline, using the STISuite V3.0 QSM GUI pipeline with Laplacian based phase unwrapping, a VSHARP background field correction and the iLSQR QSM method to calculate susceptibility maps used to estimate iron content as described previously (Li, Wu, & Liu, 2013; Spence et al., 2022).

### Plasma markers

Blood samples were taken from unfasted participants. A full blood count was carried out within NHS laboratories as described in the STRADL cohort profile (Habota, et al., 2019). CRP levels were described here as undetectable (<4 mg/L - below detection limit), normal (4-10 mg/L) or high (>10mg/L).

As described previously (Spence et al., 2024), plasma samples were assessed for sTfR (RD194011100 - Oxford Biosystems Ltd UK), ferritin (RAB0197-1KT, Merck Life Science UK Ltd) via enzyme linked immunosorbent assays (ELISA). Absorbance was read at 450nm on a uQuant BioTek microplate spectrophotometer (Biotek Instruments, Inc). Macrophage colony-stimulating factor (M-CSF; Authentikine KE00184, Proteintech Group, Inc), IL1β (BMS224-2TEN – Thermo Fisher Scientific Inc.) and IL6 (BMS213-2TEN – Thermo Fisher Scientific Inc.) were also analysed via ELISA. Absorbance was read at 450nm on an Infinite 200 PRO Tecan microplate reader.

Elemental analysis (Na, Mg, P, K, Ca, Mn, Fe, Co, Cu, Zn, As, Se, Mo, Cd, Hg and Pb) was carried out using an Agilent 7700x ICP-MS, following sample digestion with nitric acid and hydrogen peroxide.

### Environmental exposure estimation

Participant exposure to ambient atmospheric pollutants were estimated from modelled background air concentrations (DEFRA, 2023) and atmospheric emission inventories (NAEI, 2023) available for the UK. The modelled background air concentration maps provide estimates of ambient air pollutant concentrations, incorporating both local point sources and distal sources when accounting for atmospheric dispersion (Pugsley et al., 2022). Whereas the atmospheric emission inventory maps represent the total mass of pollutant released to the atmosphere within a given grid cell from all sectors, and do not account for pollutant dispersion (Tsagatakis et al., 2023). The spatially distributed time series data were both available at spatial resolutions of 1 × 1 km. The annual limits are based on the UK The Air Quality Standards Regulations 2010 (Supplementary Table 1).

### Statistical analysis and machine learning

Principal component analysis was implemented in R statistical software (v4.1.2; R Core Team 2021) with packages ‘factoExtra’, ‘tidyverse’, ‘gridExtra’, ‘Hmisc’ and ‘corrplot’ (Kassambara 2020; Wickham 2024; Auguie 2022; Harrell et al., 2019; Wei et al., 2019) to generate a *g* factor for general cognition from composite cognitive assessment scores (Mill Hill Vocabulary test, controlled word association task, logical memory subset (immediate and delayed recall), digit symbol coding task and the matrix reasoning test. Here, the *g* factor was defined as the first principal component (Figure 2A-B) and participants were divided into three classes based on tertiles of *g* factor, where class one represented those within the highest tertile of *g* factors and class three represented those within the lowest tertile of *g* factors. Demographic information regarding each tertile is detailed in Table 1. For analysis of classifiers of individual cognitive assessment scores, the median integer score was used as a cut off for high and low groups.

**Table 1.** Cohort demographics in the low, medium and high *g* factor tertiles respectively.

|  | Low | Medium | High |
| --- | --- | --- | --- |
| Age (mean ± sd) | 62.1 ± 4.9 | 61.2 ± 3.8 | 61.4 ± 5.4 |
| Sex (M:F) | 59:49 | 44:64 | 47:61 |
| Immediate Recall (mean ± sd) | 12.6 ± 2.7 | 15.2 ± 2.4 | 18.7 ± 2.2 |
| Delayed Recall (mean ± sd) | 11.2 ± 2.9 | 14.2 ± 2.7 | 17.7 ± 2.2 |
| Digit Symbol Task (mean ± sd) | 60.6 ± 12.0 | 66.8 ± 12.6 | 72.0 ± 12.5 |
| Matrix Reasoning (mean ± sd) | 6.7 ± 2.1 | 8.75 ± 1.8 | 10.0 ± 2.1 |
| Mill Hill Vocabulary Score (mean ± sd) | 29.4 ± 3.4 | 31.9 ± 3.1 | 34.8 ± 3.2 |
| Verbal Fluency (mean ± sd) | 35.4 ± 9.5 | 42.4 ± 9.9 | 49.7 ± 11.0 |

**Figure 2.**
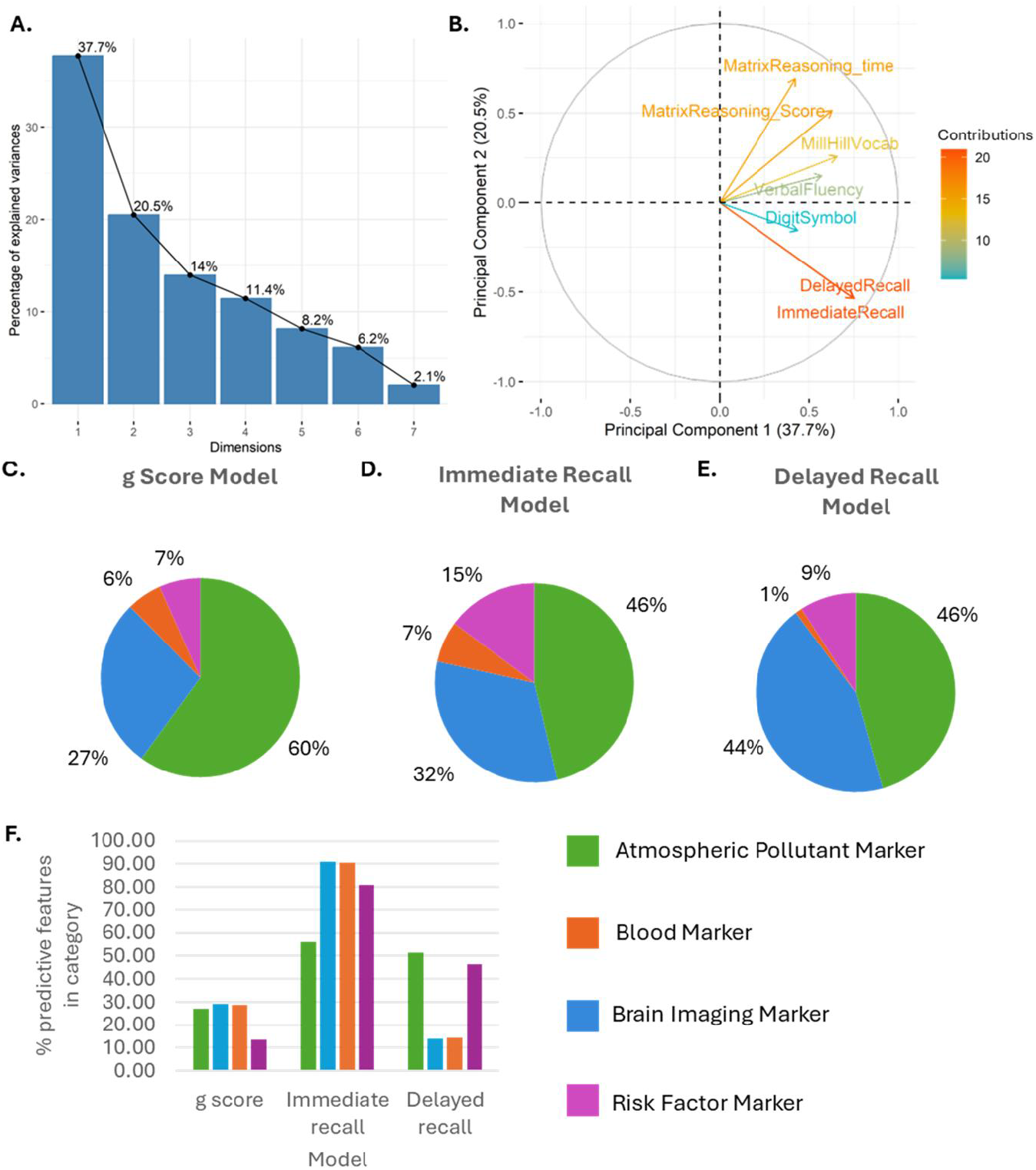
Pie charts to show percentage of predictors from each feature domain. Atmospheric pollutant markers constituted the largest domain of predictors in every model. **A)** Scree plot to show the variance explained by each principal component. General cognition (*g* factor) was defined as the first dimension (principal component 1). **B)** Biplot to show the loadings of each included cognitive variable on the first principal component. **C)** Percentage of predictors from each feature domain shown to predict *g* factor for general cognition after leave-one-out analysis. **D)** Percentage of predictors from each feature domain shown to predict immediate recall memory score after leave-one-out analysis. **E)** Percentage of predictors from each feature domain shown to predict delayed recall memory score after leave-one-out analysis. **F)** Percentage of predictive features within each category by model.

457 features were extracted from clinical, imaging, plasma analysis and environmental exposure data and these data from 80% of the cohort were used to train random forest classifiers (Supplementary Table 2; Figure 1). Classification of subjects was performed using the random forest implementation in Knime 5.2.5. (Berthold et al., 2008). Classifiers were trained with 1,000 trees per classifier and information gain as the split criterion and tested and validated on the remaining 20% of the data set. Sensitivity and specificity scores were calculated, and cross-validation was performed to assess model validity (*Tables 2-6*). Finally, leave-one-out analyses determined the predictive power of each feature (Supplementary Tables 3-5). This analysis was performed for each cognitive assessment measure. Features that were shown as strongest predictors in machine learning modelling were visualised in R statistical software (v4.1.2; R Core Team 2021) with the ggplot2 (Wickham 2016) package. Feature comparison was completed for the top predictive factors in each analysis. Shapiro Wilk’s test was initially used to determine whether data distribution was parametric. Wilcox test or Pairwise Wilcox test with Holm post-hoc correction was used to analyse non-parametric data. *T*-test or ANOVA with Tukey-HSD post-hoc correction was used to analyse parametric data.

**Table 2.**
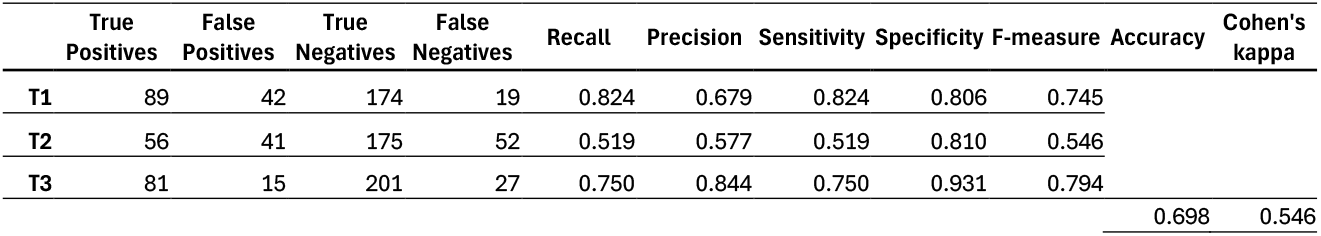
Cross-validation of the random forest model for *g* factor classification. Here, T1 is the tertile of the highest performing group and T3 is the tertile of the lowest performing group.

As mean corpuscular haemoglobin concentration and pollution measures were the only measures consistently predictive with significant differences observed in feature analysis, relationships between these two variables were explored further using linear and quadratic regression modelling employing the *lm* function in the R stats package (version 3.6.2).

## Results

### Machine learning model classifies lowest cognition score group with high sensitivity and specificity

This non-biased machine learning approach where brain imaging, blood, socio-economic and environmental exposure measures were used to model cognitive performance was able to classify general cognition tertiles with 0.70 overall accuracy. The resultant model was most sensitive and specific in classifying those with the lowest cognitive scores (75.0% and 93.1% respectively; Table 2). When cognitive assessment measures were individually assessed, random forest models were sensitive and specific at classifying low immediate recall (68.3% and 82.5% respectively, Table 3) and delayed recall (69.5% and 80.1% respectively, Table 4) scores. However, modelling with the included 443 features lacked sensitivity when classifying low digit symbol task (34.9%, Table 5) and matrix reasoning task (41.2%, Table 6) scores. Due to the reduced accuracy of the latter two models, feature analysis was not carried out for models of these cognitive test scores.

**Table 3.**
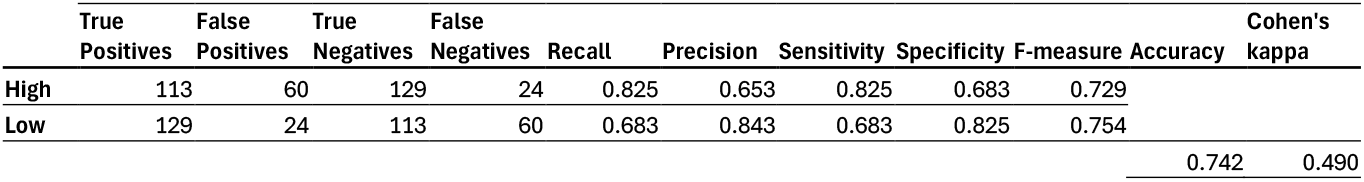
Cross-validation of the random forest model for memory score (immediate recall) classification.

**Table 4.**
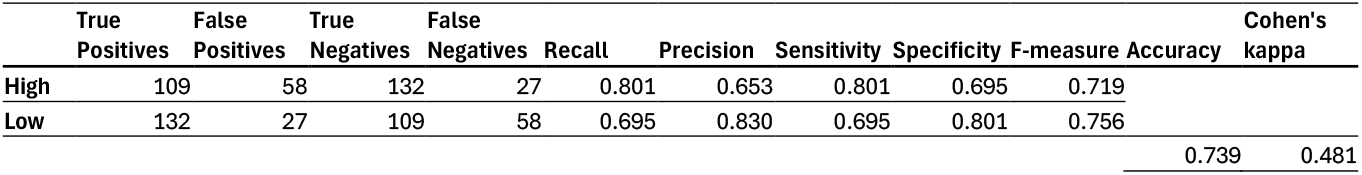
Cross-validation of the random forest model for memory score (delayed recall) classification.

**Table 5.**
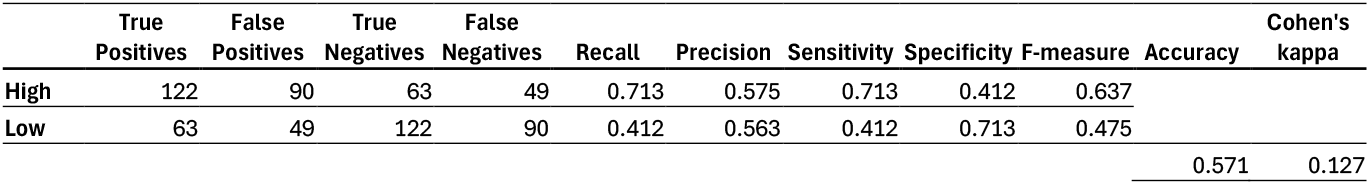
Cross-validation of the random forest model for digit symbol task score classification.

**Table 6.**
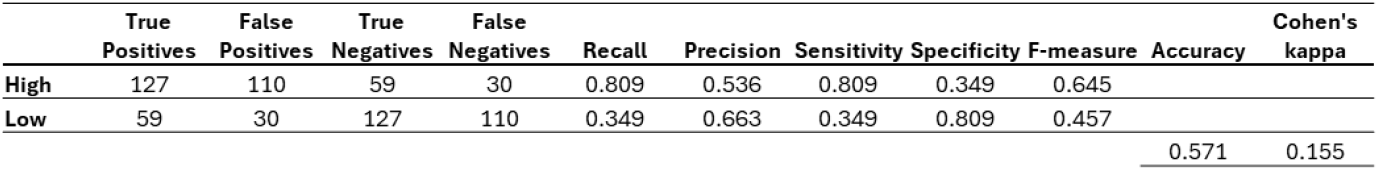
Cross-validation of the random forest model for matrix reasoning task score classification.

Brain imaging and atmospheric pollution markers represented the largest proportion of predictive markers (Figure 2C-E) although it should be noted that these two categories also had the largest number of markers contributing to the model. When normalised by the number of markers in each category, around 27-29% of atmospheric pollution, brain imaging and blood markers were predictive of general cognition (*g* factor), compared to only 13.5% of social deprivation markers (Figure 2F). More markers were predictive in general for immediate recall score with over 80% of brain imaging, blood and social deprivation markers and over 50% of atmospheric pollution markers being predictive (Figure 2F). Over 40% of atmospheric pollution and social deprivation markers were predictive for delayed recall compared to less than 15% blood and social deprivation markers (Figure 2F).

### Specific measures of education and deprivation predicted immediate recall abilities more than other aspects of cognitive performance

Whilst only 7 out of 52 social deprivation measures improved the model of general cognition score, 42 of the 52 social deprivation measures improved the model of immediate recall score whilst 24 of these measures improved the delayed recall model (Figure 2).

Measures of school attainment, rate of overcrowding and number of homes with non-central heating in the participants SIMD region improved model performance for classifying poor general cognition (Supplementary Table 3). Age, sex and education also improved general cognition model performance, however, other socioeconomic and demographic factors included in the model for classifying general cognition did not improve model performance. After post-hoc analysis there was a significantly lower education level in those with low general cognition compared to those with high general cognition (*χ*^*2*^ = 89.748, *p* <0.001; Supplementary Table 3; Figure 3A). Those with low *g* factor also lived in regions with significantly higher rate of overcrowding compared to those with high *g* factors (*p* = 0.007; Supplementary Table 3; Figure 3A). There was no significant difference between the weight of participants dependent on general cognition score tertile (Supplementary Table 3; Figure 3A).

**Figure 3.**
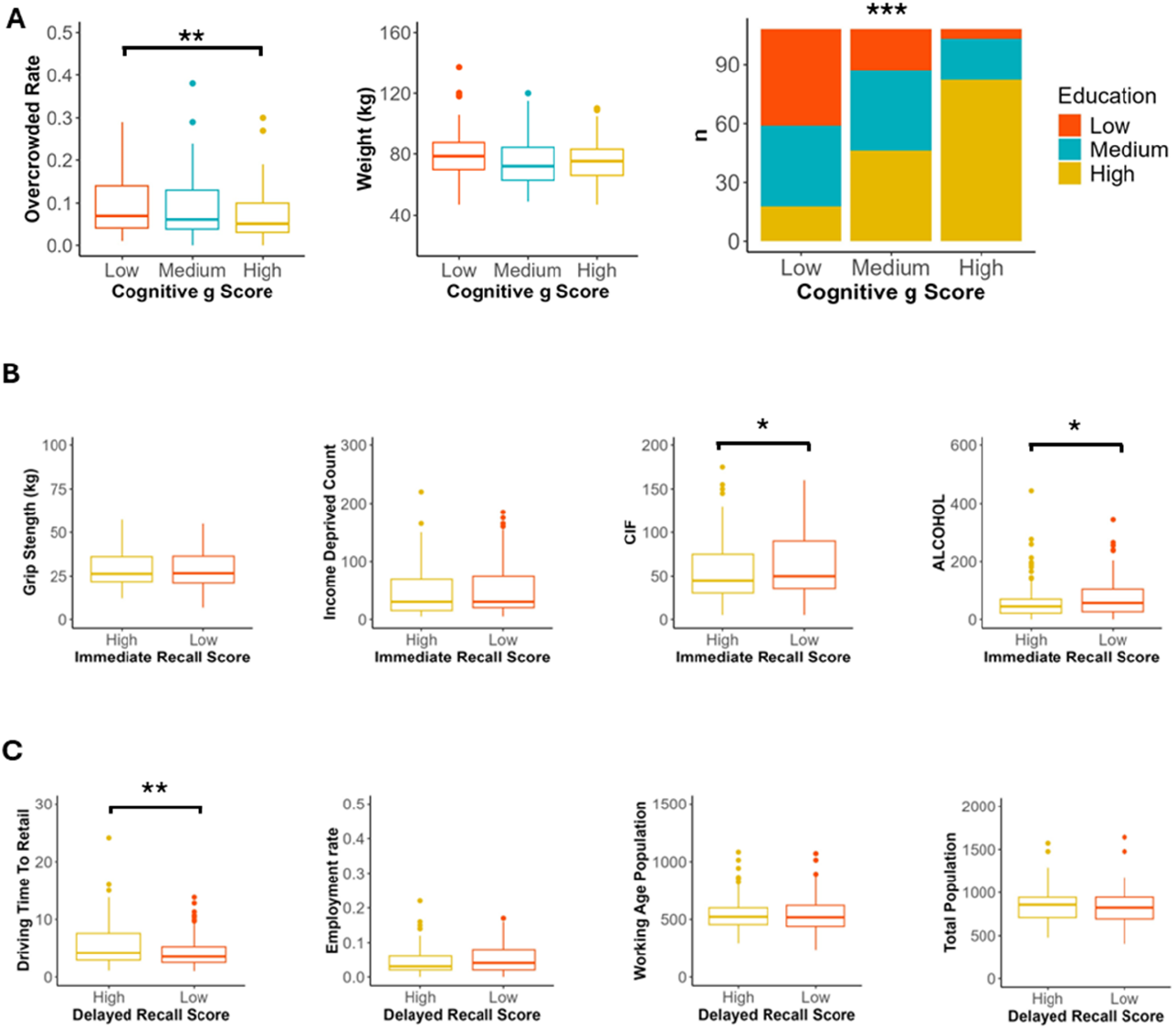
Specific measures of education and deprivation predicted immediate recall abilities more than other aspects of cognitive performance. **A)** Three socio-economic factors markers that improved the random forest classifier for *g* factor group by the greatest amount in leave-one-out analysis. Box and whisker plots demonstrate that those with lower *g* factor generally lived in regions with significantly higher rates of overcrowding (Pairwise Wilcoxon, Holm), non-significantly weighted more (Pairwise Wilcoxon, Holm) and had significantly lower levels of education (Chi Squared test). **B)** Four socio-economic factors markers that improved the random forest classifier for immediate recall score group by the greatest amount in leave-one-out analysis. Box and whisker plots demonstrate that those with lower immediate recall score had non-significantly lower grip strength and lived in regions with significantly higher comparative illness factor and hospital stays related to alcohol use and non-significantly higher income deprivation (Pairwise Wilcoxon, Holm). **C)** Four socio-economic factors markers that improved the random forest classifier for immediate recall score group by the greatest amount in leave-one-out analysis. Box and whisker plots demonstrate that those with lower immediate recall score lived in regions with significantly less driving travel time to retail, non-significantly higher percentage of people who are employment deprived, and non-significantly lower working age population and total population estimate (Pairwise Wilcoxon, Holm). \**p* <0.05, \*\**p* <0.01, \*\*\**p* <0.001

In the immediate recall model, grip strength, income deprivation, comparative illness factor and emergency hospital visits resulting from alcohol were the strongest predictors. Feature comparison exhibited significantly higher comparative illness factor and number of emergency hospital stays due to alcohol in the low compared to high immediate recall score group (*p* = 0.046 and *p* = 0.023 respectively; Supplementary Table 4; Figure 3B). There was no significant difference between grip strength or income deprivation between low and high immediate recall performers.

The strongest social-deprivation predictors for delayed recall score were driving time to retail, employment rate, working age population and total population. Feature comparison demonstrated that those with low scores in the delayed recall task lived in areas with significantly shorter driving time to get to a retail area (*p* = 0.001; Supplementary Table 5; Figure 3C). There was no significant difference between employment rate, working age or total population between delayed recall score groups.

### Atrophy, swelling and iron content of specific brain regions predicted poorer cognitive performance

Whilst 29 out of 116 imaging measures improved the model of general cognition score, 91 of the 116 imaging improved the model of immediate recall score whilst 16 of these measures improved the delayed recall model (Figure 2).

Volumes of the right cerebellum (cortex), corpus callossum and left postcentral cortex and iron in the right amygdala improved model classification performance for poor general cognition (Supplementary Table 3; Figure 4A). There were no significant differences in these imaging measures between *g* factor groups. In the immediate recall model, right caudal anterior cingulate cortex, right Bankssts cortex, left insula and right isthmus cingulate cortex volumes were the strongest predictors. Feature comparison exhibited significantly lower left insula cortex in the low compared to high immediate recall score group (*p* = 0.025; Supplementary Table 4; Figure 4B). There was no significant difference between other volumes in low compared to high immediate recall performers. The strongest imaging predictors for delayed recall score were iron in the left amygdala and right thalamus and volume in the right cerebral white matter and left ventral diencephalon. Feature comparison demonstrated no significant differences between these measures in the low delayed recall group compared to the high delayed recall group (Supplementary Table 5; Figure 4C).

**Figure 4.**
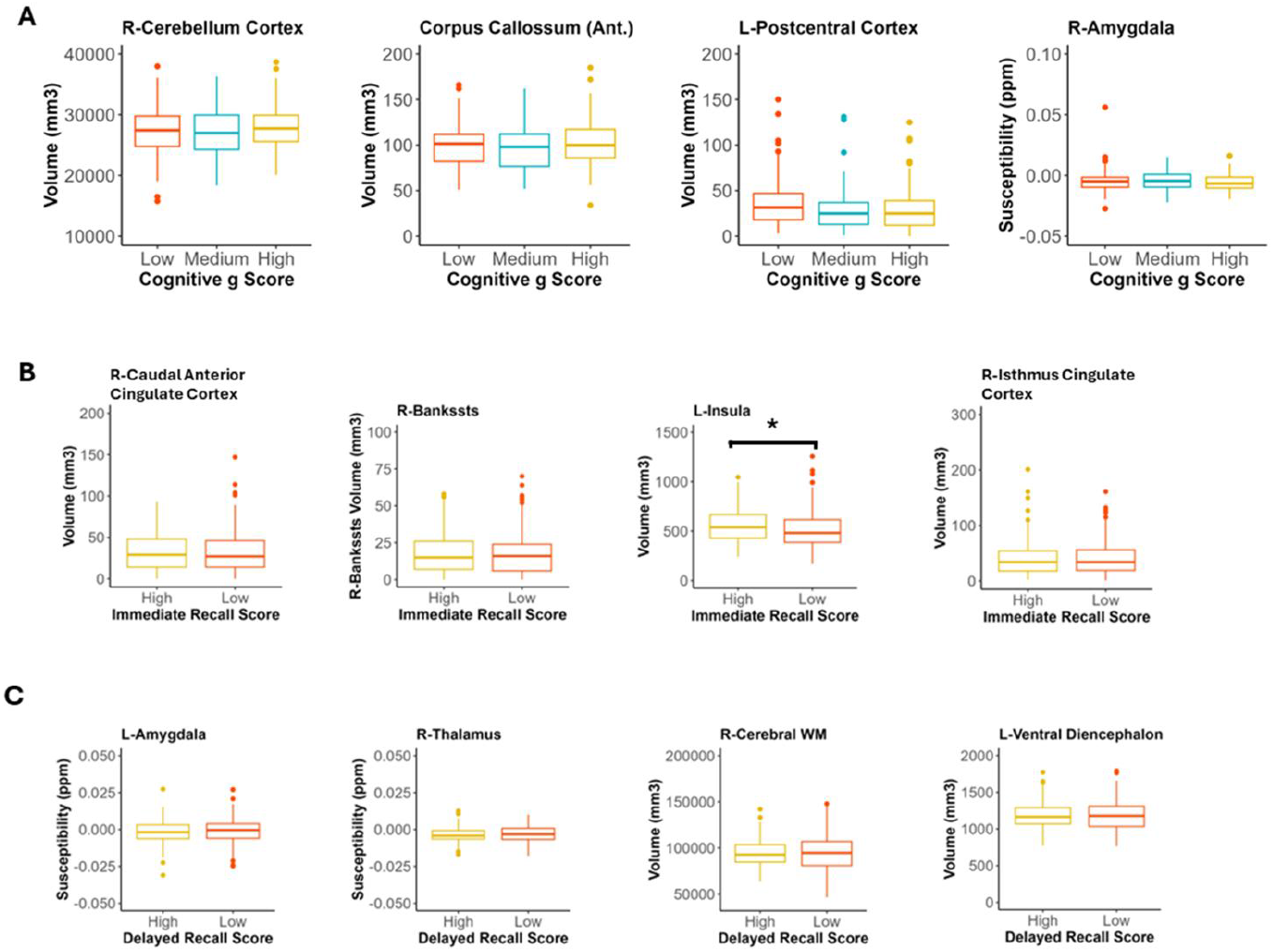
Atrophy, swelling and iron content of specific brain regions predicted poorer cognitive performance. **A)** Four brain imaging measures that improved the random forest classifier for *g* factor group by the greatest amount in leave-one-out analysis. Box and whisker plots demonstrate that those with lower *g* factor generally had non-significantly lower volumes of large regions such as the right cerebellar cortex and anterior corpus callossum, but non-significantly higher volume of the left postcentral cortex. Those with lower *g* factors also had non-significantly higher levels of iron in the right amygdala (Pairwise Wilcoxon, Holm). **B)** Four brain imaging measures that improved the random forest classifier for immediate recall score group by the greatest amount in leave-one-out analysis. Box and whisker plots demonstrate that those with lower immediate recall score non-significantly higher volumes of the right caudal anterior cingulate cortex but significantly lower insula volume and non-significantly lower volume of the bankssts cortex and right isthmus cingulate cortex (Pairwise Wilcoxon, Holm). **C)** Four brain imaging measures that improved the random forest classifier for immediate recall score group by the greatest amount in leave-one-out analysis. Box and whisker plots demonstrate that those with lower delayed recall score exhibit non-significantly higher levels of iron in the left amygdala (Pairwise Wilcoxon, Holm) and right thalamus (ANOVA, Tukey-HSD) and non-significantly lower volumes of the left ventral diencephalon (Pairwise Wilcoxon, Holm) and right cerebral white matter (ANOVA, Tukey-HSD). \**p* <0.05, \*\**p* <0.01, \*\*\**p* <0.001

### Blood measures of anaemia and chronic inflammation consistently predicted poorer cognitive performance

Out of the 21 blood markers assessed, 6 improved the model of general cognition score, 19 improved the model of immediate recall score whilst only 3 improved the delayed recall model. Measures of mean corpuscular haemoglobin and CRP were predictors across each of the three models (Figure 2).

Measures of haematocrit, haemoglobin, mean corpuscular haemoglobin and mean corpuscular volume each improved model performance for classifying poor general cognition (Supplementary Table 3; Figure 5A). In the immediate recall model, mean corpuscular haemoglobin, white blood cell count, ferritin and zinc levels were the strongest predictors (Supplementary Table 4; Figure 5B). The strongest blood marker predictors for delayed recall score were mean corpuscular haemoglobin, CRP and basophil count (Supplementary Table 5; Figure 5C). Feature comparison demonstrated no significant differences between blood markers by *g* factor, immediate recall or delayed recall group.

**Figure 5.**
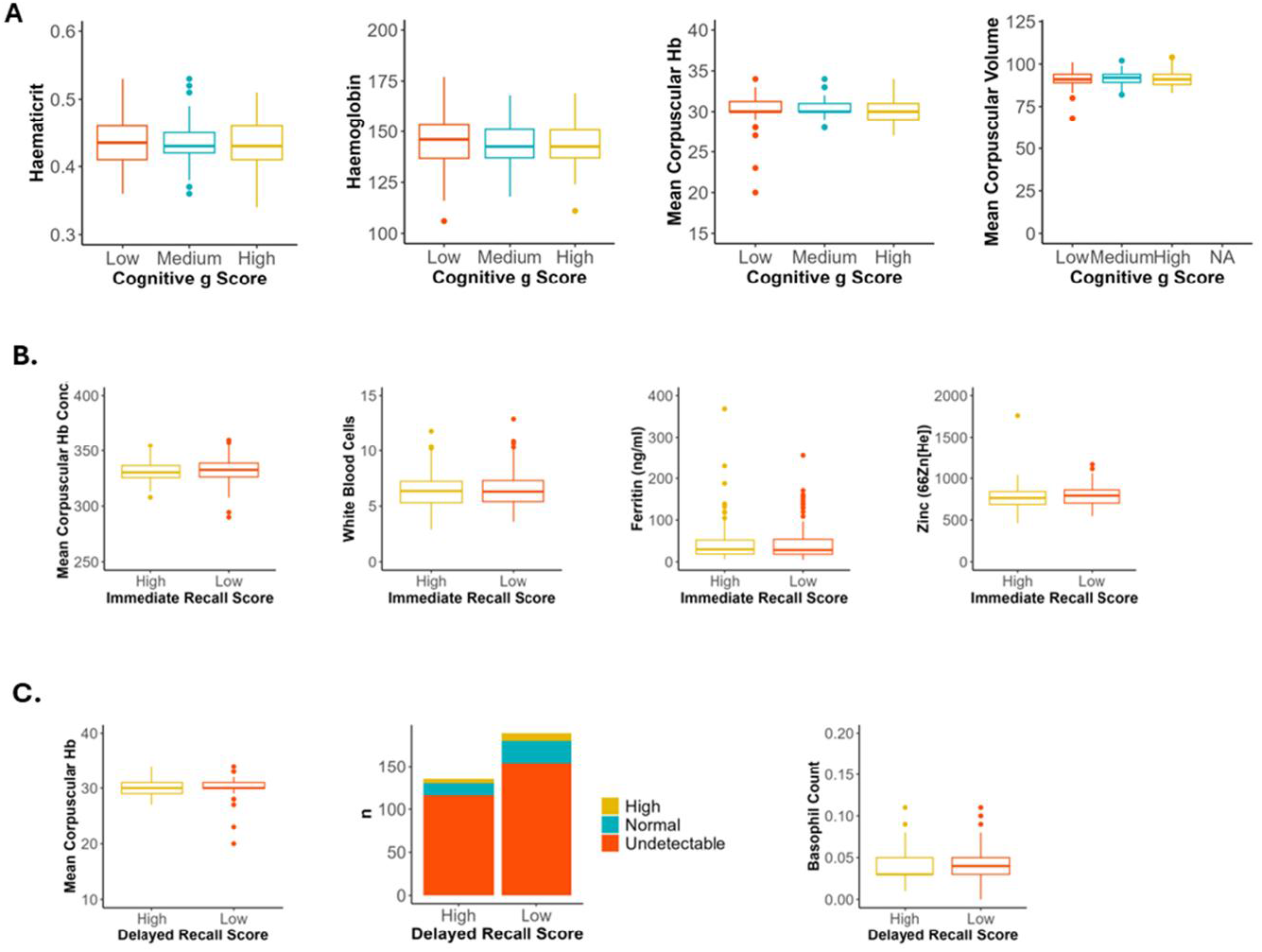
Blood measures of anaemia and chronic inflammation consistently predicted poorer cognitive performance. **A)** Four blood markers that improved the random forest classifier for *g* factor group by the greatest amount in leave-one-out analysis. Box and whisker plots demonstrate that those with lower *g* factor had non-significantly higher haemoglobin, haematocrit and mean corpuscular haemoglobin, but non-significantly lower mean corpuscular volume (Pairwise Wilcoxon, Holm**). B)** Four blood markers that improved the random forest classifier for immediate recall performance by the greatest amount in leave-one-out analysis. Box and whisker plots demonstrate that those with lower immediate recall score had non-significantly higher mean corpuscular haemoglobin concentration, white blood cell count and plasma Zinc but non-significantly lower plasma ferritin (Pairwise Wilcoxon, Holm). **C)** Three blood markers that improved the random forest classifier for delayed recall performance in leave-one-out analysis. Box and whisker plots demonstrate that those with lower delayed recall score had non-significantly higher mean corpuscular haemoglobin, basophil count (Pairwise Wilcoxon, Holm) and C-reactive protein (Chi squared test). \**p* <0.05, \*\**p* <0.01, \*\*\**p* <0.001

### Environmental exposure to heavy metals, PM25, SO_2_ and CO were consistently the strongest predictors of poor cognitive performance

Across all three assessed models, atmospheric pollutant exposure was consistently the strongest predictor of cognitive performance. Of 254 measures of atmospheric pollutant levels in the region of dwelling, 63 measures improved the model of general cognition score, 131 measures improved the model of immediate recall score whilst 120 measures improved the delayed recall model (Figure 2). Upon feature analysis, measures of Arsenic, cadmium, copper, lead, mercury and PM2.5 were consistently identified as predictors of poorer *g* factor, immediate recall score and delayed recall score.

Levels of arsenic, carbon monoxide, lead and PM2.5 in the years of measurement closes to those of cognitive testing improved model classification performance for poor general cognition. Significantly lower levels of arsenic (2015), carbon monoxide (2010), lead (2016) and PM2.5 (2017) were observed in the high *g* factor group compared to the medium *g* factor group (*p* = 0.013, *p* = 0.005, *p* = 0.003, *p* = 0.008 respectively; Figure 6A). Significantly higher levels of carbon monoxide (2010), lead (2016) and PM2.5 (2017) were observed in the low *g* factor group compared to the high *g* factor group (*p* = 0.011, *p* = 0.003, *p* = 0.009 respectively; Supplementary Table 3; Figure 6A).

**Figure 6.**
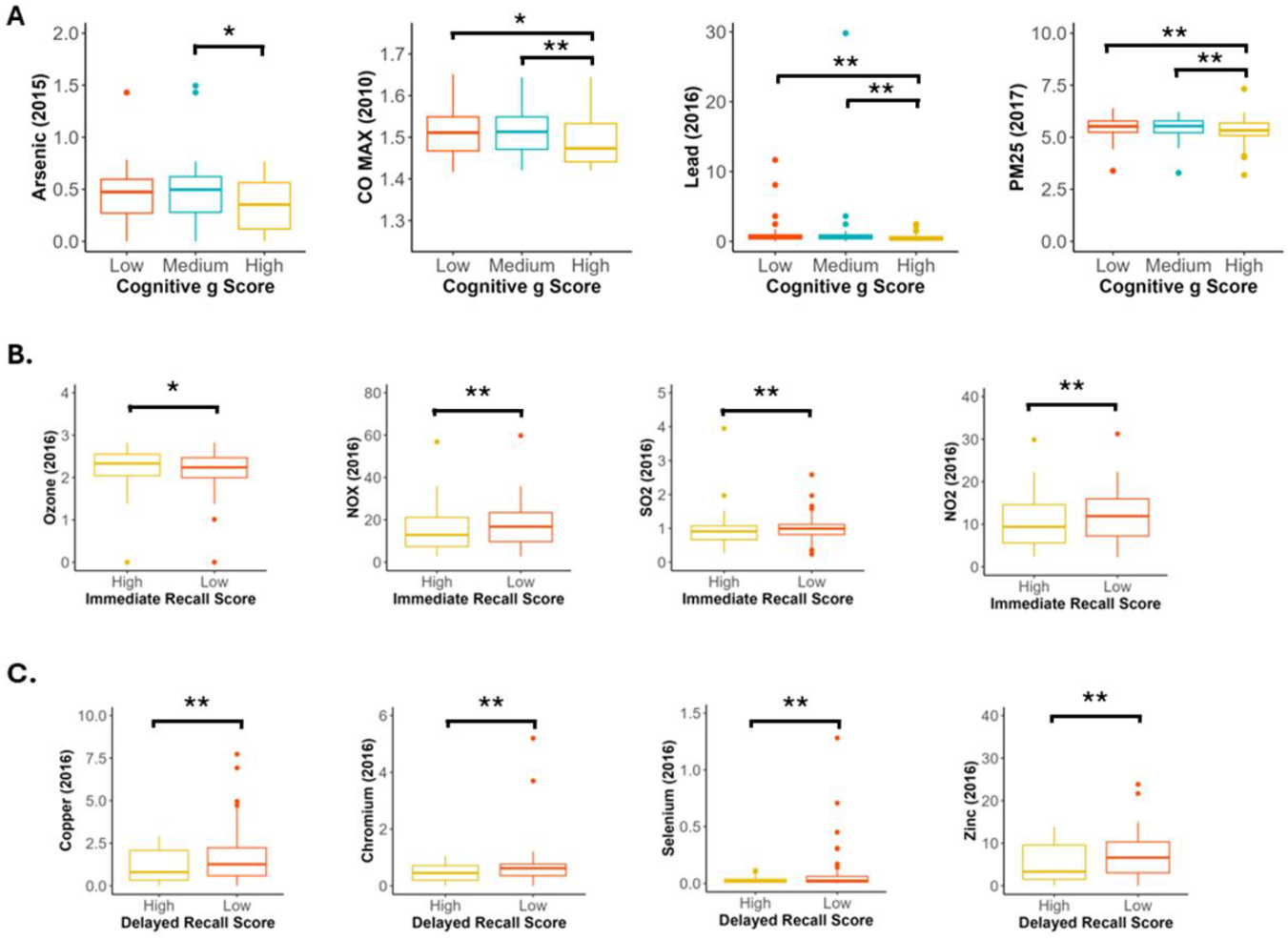
Environmental exposure to heavy metals, PM25, SO2 and CO were consistently the strongest predictors of poor cognitive performance. **A)** Four modelled air quality and emissions measures that improved the random forest classifier for *g* factor group by the greatest amount in leave-one-out analysis. Box and whisker plots demonstrate that those with lower *g* factor lived in regions with significantly higher levels of Arsenic, carbon monoxide, lead and particulate matter in the year of measurement closest to the year of cognitive assessment (Pairwise Wilcoxon, Holm). **B)** Four modelled air quality and emissions measures that improved the random forest classifier for immediate recall score by the greatest amount in leave-one-out analysis. Box and whisker plots demonstrate that those with lower immediate recall score lived in regions with significantl higher levels of NOX, NO2 and SO2 but significantly lower levels of Ozone in the year of measurement closest to the year of cognitive assessment (Pairwise Wilcoxon, Holm). **C)** Four modelled air quality and emissions measures that improved the random forest classifier for immediate recall score by the greatest amount in leave-one-out analysis. Box and whisker plots demonstrate that those with lower delayed recall score lived in regions with significantly higher levels of copper, chromium, selenium and zinc in the year of measurement closest to the year of cognitive assessment (Pairwise Wilcoxon, Holm). \**p* <0.05, \*\**p* <0.01, \*\*\**p* <0.001

In the immediate recall model, ozone, oxides of nitrogen, nitrogen dioxide and sulphur dioxide were the strongest predictors. Feature comparison revealed significantly lower levels of ozone in the low compared to high immediate recall group (*p* = 0.043; Figure 6B). There were significantly higher levels of NOX, NO2 and SO2 in the low immediate recall score group compared to the high immediate recall score group (*p* = 0.008, *p* = 0.008, *p* = 0.006 respectively; Supplementary Table 4; Figure 6B).

The strongest imaging predictors for delayed recall score were copper, chromium, selenium and zinc. Feature comparison demonstrated higher levels of copper, chromium, selenium and zinc in the low delayed recall group compared to the high delayed recall group (*p* = 0.001, *p* = 0.008, *p* = 0.006, *p* = 0.001 respectively; Supplementary Table 5; Figure 6C).

### Low mean corpuscular haemoglobin concentration is a biomarker of pollution exposure and poor cognition

As mean corpuscular haemoglobin concentration (MCHC) and levels of arsenic, cadmium, copper, lead, mercury and PM25 in the region of dwelling were key predictive features across all three models of cognitive health, the relationships between these markers and general cognition were assessed. Whilst MCHC was not significantly associated with general cognition score, there was a significant quadratic relationship between MCHC and immediate recall memory score (Figure 7A; Supplementary Figure 1). Low MCHC was also significantly correlated with increased exposure with increased dwelling region atmospheric levels of arsenic (*ρ* = −0.132, *p* = 0.018; Figure7B), cadmium (*ρ* = −0.200, *p* <0.001; Figure7C), copper (*ρ* = −0.154, *p* = 0.006; Figure7D), lead (log transformed, *ρ* = −0.131, *p* = 0.019; Figure7E), mercury (log transformed, *ρ* = −0.118, *p* = 0.034; Figure7F) and PM25 (*ρ* = −0.164, *p* = 0.003; Figure7G). Our results show that in areas with higher levels of atmospheric pollution individuals had generally lower MCHC (Figure 7) and generally lower memory score (Figure 6). MCHC could therefore represent a biomarker for pollution-related poor memory function.

**Figure 7.**
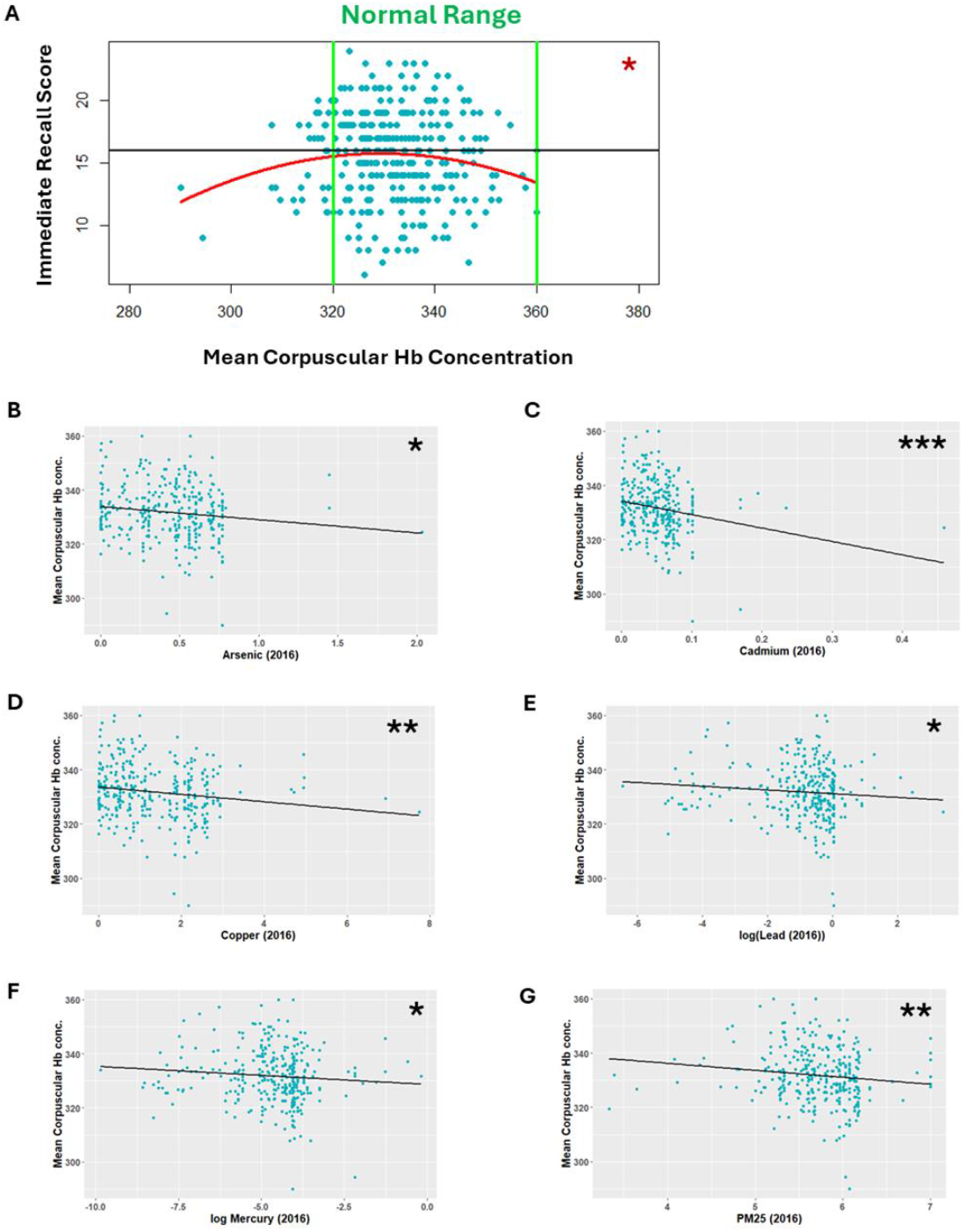
Clinically low mean corpuscular haemoglobin concentration is associated with lower immediate recall score and increased levels of predictive pollutants. **A)** Mean corpuscular haemoglobin demonstrates a quadratic relationship with immediate recall score (*p* = 0.0253). In the year of measurement that was predictive closest to the years of cognitive assessment lower mean corpuscular haemoglobin was significantly associated with increased dwelling region atmospheric levels of **B)** Arsenic (*p* = 0.018, *ρ* = −0.132), **C)** Cadmium (*p* <0.001, *ρ* = −0.200), **D)** Copper (*p* = 0.006, *ρ* = −0.154), **E)** lead (log transformed, *p* = 0.019, *ρ* = −0.131), **F)** Mercury (log transformed, *p* = 0.034, *ρ* = −0.118), **G)** PM25 (*p* = 0.003, *ρ* = −0.164). \**p* <0.05, \*\**p* <0.01, \*\*\**p* <0.001

## Discussion

Here, we use a non-biased machine learning approach to identify key predictors of cognitive performance in a unique birth cohort with detailed demographic information. We show differences in predictive measures for different cognitive domains, where the highest number of predictors were identified for immediate recall scores. This indicates differing importance of modifiable risk factors for each cognitive domain, with measures of combustion emissions and anaemia representing the features which most improved the model of general cognition, specific imaging markers improving the immediate recall model by the greatest amount and atmospheric pollutants indicative of smelting and pesticide use most improved the delayed recall model. Importantly, we demonstrate that low mean corpuscular haemoglobin levels were significantly increased measures of atmospheric arsenic, cadmium, copper, lead, mercury and PM2.5 in the region of dwelling, all of which are associated with poorer memory performance. Mean corpuscular haemoglobin concentration therefore represents a potential routinely assessed biomarker for risk of dementia in those living in areas of high pollution.

### Poor geographic, health, education and income access increase dementia risk

Measures indicative of geographic access, health and income were specifically predictive of memory scores. This suggests a heavier impact of social deprivation on memory aspects of cognition. In the general cognition model, only 3 measures of social deprivation were predictive of poorer *g* factor, however, 41 social-deprivation measures were predictive of specific memory scores. As immediate recall tasks have shown greatest sensitivity of predicting dementia (Fayemiwo et al., 2023), this result indicates that social deprivation is a key risk factor for dementia. The three measures consistent across models (school attainment, overcrowding and number of homes with non-central heating) highlight that education and poverty indicators likely play a key role in determining general cognitive health.

### Atrophy and swelling detected on brain MRI are identifiable risk factors for poor cognitive function

There were also differences in the imaging markers predictive of general cognition compared to specific measures of memory performance. Whilst there were consistencies across models in the links between the volumes of regions involved in connectivity and signal relay such as the corpus callossum and thalamus, there were some regions such as the temporal cortex and inferior parietal cortex which were only predictive in classifying immediate recall memory score, regions that have been previously shown to play specific roles in memory function (Leff et al., 2009; Cabeza et al., 2009). Atrophy of large regions such as the cerebellum was shown to predict general cognitive performance as well as specific memory scores. As these large regions can be visualised at a lower resolution, this opens opportunities for monitoring with low field MRI which is less expensive and more portable than conventional 3T and above MRI, with implications for improving health equity.

Further, whilst atrophy of most regions was associated with poorer cognitive performance, consistent with previous studies (Saykin et al., 2006; Jack et al., 2005; McEvoy et al., 2009), some regions such as the inferior temporal cortex demonstrated association between increased volume and poorer cognitive performance. Importantly, this suggests that swelling, rather than atrophy, of some regions may be associated with poorer cognitive performance. This local swelling could be caused by inflammation of the myelin sheath which has been shown to occur during traumatic brain injury leading to local swelling effects and accumulation of proteins (Sabharwal & Koushika, 2019).

### Living in regions with high vehicle emissions, refuse incineration and fossil fuel combustion may increase risk of poor cognitive health

Increased measures of atmospheric arsenic, cadmium, copper, lead, mercury and PM2.5 in the region of dwelling are associated with poorer cognitive performance. This association holds for measurement of pollutants not only in the years closest to cognitive testing but also those measured years before cognitive testing. This is consistent with previous studies that have demonstrated a 4–10-year lag in the association between PM2.5 and memory decline (Tonne et al., 2014; Chen et al., 2017). As demonstrated by Jung et al. (2015), there may be a threshold of certain pollutants that must be met before an effect on cognitive health can be observed. This may explain why pollutants were not predictive of cognitive decline in some years. Arsenic, cadmium, lead and mercury have previously been identified as significant to public health and increased atmospheric levels of these four specific heavy metals in the UK has been linked with a rise in refuse incineration, steel production, fossil fuel combustion (Hutton & Symon, 1986; Tchounwou et al., 2012). Some studies suggest that a combination of arsenic, cadmium and lead may increase neurotoxicity through combined alteration of the ERK-JNK pathway, promoting apoptosis (Rai et al., 2010). It may be therefore, that the threshold for each of these exposures is lower than other heavy metals. Due to the charged nature of heavy metals including copper, they are able to strongly bind to particulate matter and are in turn known to increase reactive oxygen species and oxidative stress which may explain their associations with cognitive performance observed in this study (Tacu, Kokalari et al. 2021).

### Routinely assessed measures of anaemia represent an accurate predictive measure of pollution-dependent poor cognitive function

We show that MCHC consistently predicted poorer cognitive health across all three assessed models and after feature analysis it was determined that MCHC demonstrated a significant quadratic relationship with immediate recall memory score. This finding corroborates previous studies that show both low and high levels of haemoglobin as associated with dementia in a u-shaped association, whilst anaemia was associated with increased risk of dementia (Wolters et al., 2019; Shah et al., 2008) and highlights the potential for identifying at risk groups through routine in clinic blood testing. It is thought that higher MHCH is indicative of microcytic anaemia and/or polycythaemia often caused by iron deficiency or excessive alcohol consumption/smoking respectively, are detrimental to cognitive health likely due to impaired oxygen carrying capacity (Pearson, 1991). However, low MCHC is associated with iron-deficiency anaemia and we show that lower MCHC is significantly associated with increased levels of dwelling region atmospheric pollutant levels. This fits with previous findings that suggest PM25 and cadmium exposure in particular can induce iron-deficiency anaemia (He et al., 2023; Li et al., 2024; Hwang et al., 2024; Fujiwara et al., 2020). Our results highlight the potential for MCHC as a routinely assessed biomarker of pollution exposure related cognitive risk.

Whilst heavy metal exposure (e.g. atmospheric lead) was a strong predictor of poor cognitive performance, blood levels of such heavy metals were not predictive of general cognition. It is likely that heavy metal exposure results in end organ accumulation of these heavy metals and so blood levels were not reflective of total body levels (Jaishankar et al., 2014). However, blood markers of heavy metals were predictive of memory scores, suggesting that this may be easier to track in those at risk of Alzheimer’s type dementia. Atmospheric pollutant measures could be utilised to identify at risk groups and “green corridors” of low levels of predictive pollutants to inform public footpaths and walk-to-school or -work routes. Recent studies have shown promise in combatting the effects of exposure to pollutants such as arsenic (Roy et al., 2006). Therefore, if at risk groups are identified through atmospheric pollutant mapping, there is opportunity for targeted preventative treatment strategies.

It is important to consider the context of the studied cohort when interpreting these results. Whilst this cohort offered a fairly homogeneous foundation population for determining exposure influence, there exists an inequality in the dementia incidence across racial and ethnic groups which could not be assessed due to the lack of diversity within this regional birth cohort (Mayeda et al., 2016). It should additionally be taken into account that non-neurological diseases such as Diabetes mellitus may confound an individual’s risk for dementia, however, such disease status was not included in this study due to the lack of data on non-neurological disease status in this cohort (Xue et al., 2019).

## Conclusions

This study highlights the importance of modifiable risk factors for poor cognition which could be addressed through targeted prevention strategies and relatively inexpensive clinical implementations. Across models for general cognitive performance, immediate and delayed recall, levels of mean corpuscular haemoglobin and atmospheric pollutants including PM2.5, arsenic and lead in the year of measurement and in years prior to measurement were the strongest predictors of cognitive performance. This indicates that increased exposure to these pollutants may pose a significant detriment to cognitive health and that routinely measures MCHC may be used as a biomarker for pollution exposure related risk to cognitive health. Further longitudinal study is vital to understand the mechanisms of these factors in affecting brain health, as well as to understand the potential ways to identify and protect at risk populations on a local, national and international scale.

## Acknowledgments

We are grateful to the Aberdeen Children of the 1950’s (ACONF) subset of Generation Scotland GS:SFHS who took part in the STRADL study, supported and funded by the Wellcome Trust Strategic Award ‘Stratifying Resilience and Depression Longitudinally’ (STRADL) [104036/Z/14/Z]. Generation Scotland received core support from the Chief Scientist Office of the Scottish Government Health Directorates [CZD/16/6] and the Scottish Funding Council [HR03006] and is currently supported by the Wellcome Trust [216767/Z/19/Z]. HS is supported by the Roland Sutton Academic Trust [0076/R/19]. We also thank the STRADL project team.

## Funding

NIH (R01NS127186)

Wellcome Trust Strategic Award ‘Stratifying Resilience and Depression Longitudinally’ (STRADL) [104036/Z/14/Z].

Target ALS (BB-2022-C4-L2)

## Ethics Statement

All components of STRADL received formal, national ethical approval from the National Health Service (NHS) Tayside committee on research ethics (reference 14/SS/0039).

## Author Contribution Statement

HS, HJ, JL, JMG and FMW contributed to the conception and design of the study. GW and HS performed the MRI data analysis. HJ performed pollution data spatial extraction. GC performed ICP-MS analysis. HS and JL performed the machine learning analysis. HS and HJ prepared the draft manuscript. All authors contributed to manuscript revision, and have read and approved the final version.

## References

Auguie B. (2022). gridExtra: Miscellaneous Functions for “Grid” Graphics. R package version 2.3, https://cindyfang70.github.io/gridExtra/.

Berthold MR, Cebron N, Dill F, et al. (2008) KNIME: the Konstanz information miner. In Data Analysis, Machine Learning and Applications, Preisach C, Burkhardt H, Schmidt-Thieme L, et al. (eds). Springer: Berlin, Heidelberg, 319–326.

Cabeza R, Ciaramelli E, Olson IR, Moscovitch M. (2008) The parietal cortex and episodic memory: an attentional account. Nat Rev Neurosci. 9(8):613–25. doi: 10.1038/nrn2459.

Carey IM, Anderson HR, Atkinson RW, et al. (2018) Are noise and air pollution related to the incidence of dementia? A cohort study in London, England. BMJ Open 8:e022404. doi: 10.1136/bmjopen-2018-022404

DEFRA. (2023) Modelled background pollution data. Available online: https://uk-air.defra.gov.uk/data/laqm-background-home (accessed on 28 December 2024). © Crown copyright 2021 Defra via uk-air.defra.gov.uk, licensed under the Open Government Licence.

Dintica CS, Yaffe K. (2022) Epidemiology and Risk Factors for Dementia. Psychiatric Clinics of North America. 45(4): 677–689. doi: 10.1016/j.psc.2022.07.011.

Fayemiwo MA, Olowookere TA, Olaniyan OO, et al. (2023) Immediate word recall in cognitive assessment can predict dementia using machine learning techniques. Alzheimers Res 15(1):111. doi: 10.1186/s13195-023-01250-5.

Foubert-Samier A, Catheline G, Amieva H, et al. (2012) Education, occupation, leisure activities, and brain reserve: a population-based study. Neurobiol Aging. 33(2): 423.e15–25. doi: 10.1016/j.neurobiolaging.2010.09.023.

Fujiwara Y, Lee JY, Banno H, et al. (2020) Cadmium induces iron deficiency anemia through the suppression of iron transport in the duodenum. Toxicol Lett. 10(332):130–139. doi: 10.1016/j.toxlet.2020.07.005.

Habota T, Sandu AL, Waiter GD, et al. (2021) Cohort profile for the STratifying Resilience and Depression Longitudinally (STRADL) study: A depression-focused investigation of Generation Scotland, using detailed clinical, cognitive, and neuroimaging assessments. Wellcome Open Res. 16(4): 185. doi: 10.12688/wellcomeopenres.15538.2.

Harrell Jr F, & Dupont Ch. (2019). Hmisc: Harrell Miscellaneous. R Package Version 4.2-0. https://CRAN.R-project.org/package=Hmisc

He C, Xie L, Gu L, et al. (2023) Anemia is associated with long-term exposure to PM_2.5_ and its components: a large population-based study in Southwest China. Ther Adv Hematol. 28(14)20406207231189922. doi: 10.1177/20406207231189922.

Hong CH, Falvey C, Harris TB, et al. (2013) Anemia and risk of dementia in older adults: findings from the Health ABC study. Neurology. 81(6): 528–33. doi: 10.1212/WNL.0b013e31829e701d.

Hutton M, Symon C. (1986) The quantities of cadmium, lead, mercury and arsenic entering the U.K. environment from human activities. Science of The Total Environment. 57: 129–150 doi: 10.1016/0048-9697(86)90018-5.

Hwang J, Kim HJ. (2024) Association of ambient air pollution with hemoglobin levels and anemia in the general population of Korean adults. BMC Public Health 24, 988 doi: 10.1186/s12889-024-18492-z

Jack CR Jr, Shiung MM, Weigand SD, et al. (2005) Brain atrophy rates predict subsequent clinical conversion in normal elderly and amnestic MCI. Neurology. 65(8): 1227–31. doi: 10.1212/01.wnl.0000180958.22678.91.

Jaishankar M, Tseten T, Anbalagan N, et al. (2014) Toxicity, mechanism and health effects of some heavy metals. Interdiscip Toxicol. 7(2): 60–72. doi: 10.2478/intox-2014-0009.

Jia L, Du Y, Chu L, et al. (2020) Prevalence, risk factors, and management of dementia and mild cognitive impairment in adults aged 60 years or older in China: a cross-sectional study. The Lancet Public Health. 5(12): e661–e671. doi: 10.1016/S2468-2667(20)30185-7

Jung CR, Lin YT, Hwang BF. (2015) Ozone, particulate matter, and newly diagnosed Alzheimer’s disease: A population-based cohort study in Taiwan. J Alzheimers Dis 44:573–584. doi: 10.3233/JAD-140855

Kassambara A & Mundt F. (2020) factoextra: Extract and Visualize the Results of Multivariate Data Analyses. R package version 1.0.7.999, https://github.com/kassambara/factoextra.

Kravitz BA, Corrada MM, Kawas CH. (2009) Elevated C-reactive protein levels are associated with prevalent dementia in the oldest-old. Alzheimers Dement. 5(4): 318–23. doi: 10.1016/j.jalz.2009.04.1230.

Leff AP, Schofield TM, Crinion JT, et al. The left superior temporal gyrus is a shared substrate for auditory short-term memory and speech comprehension: evidence from 210 patients with stroke. Brain. 132(Pt 12):3401–10. doi: 10.1093/brain/awp273.

Li W, Wu B, & Liu C. (2013). 5223 STI suite: A software package for quantitative susceptibility imaging.

Li L, Ran Y, Zhuang Y, et al. (2024) Risk analysis of air pollutants and types of anemia: a UK Biobank prospective cohort study. Int J Biometeorol. 68(7):1343–1356. doi: 10.1007/s00484-024-02670-0.

Livingston G, Huntley J, Sommerlad A et al. (2023) Dementia prevention, intervention, and care: 2020 report of the Lancet Commission. The Lancet. 396(10248): 413 – 446

Mayeda ER, Glymour MM, Quesenberry CP, et al. (2016) Inequalities in dementia incidence between six racial and ethnic groups over 14 years. Alzheimers Dement. 12(3): 216–24. doi: 10.1016/j.jalz.2015.12.007.

McEvoy LK, Fennema-Notestine C, Roddey JC, et al. (2009) Alzheimer disease: quantitative structural neuroimaging for detection and prediction of clinical and structural changes in mild cognitive impairment. Radiology. 251(1): 195–205. doi: 10.1148/radiol.2511080924.

NAEI (National Atmospheric Emissions Inventory) (2023) UK emissions data gridded emissions data selector. Available online: https://naei.energysecurity.gov.uk/data/maps/download-gridded-emissions (accessed on 18 December 2024)

Pearson TC. (1991) Apparent polycythaemia. Blood Rev. 5(4): 205–13. doi: 10.1016/0268-960x(91)90010-a.

Peters R, Ee N, Peters J, et al. (2019) Air Pollution and Dementia: A Systematic Review. J Alzheimers Dis. 70(1):S145–S163. doi: 10.3233/JAD-180631.

Pugsley KL, Stedman JR, Brookes DM et al. (2022) Technical Report on UK Supplementary Modelling Assessment Under the Air Quality Standards Regulations 2010 for 2020. London, UK: Department for Environment, Food and Rural Affairs. https://uk-air.defra.gov.uk/library/reports?report_id=1022 [accessed 18 Dec 2024].

R Core Team (2021). R: A language and environment for statistical computing. R Foundation for Statistical Computing, Vienna, Austria. URL https://www.R-project.org/

Rai A, Maurya S, Khare P, et al. (2010) Characterization of Developmental Neurotoxicity of As, Cd, and Pb Mixture: Synergistic Action of Metal Mixture in Glial and Neuronal Functions, Toxicological Sciences, 118(2): 586–601, doi: 10.1093/toxsci/kfq266

Roy S, Chattoraj A, Bhattacharya S. (2006) Arsenic-induced changes in optic tectal histoarchitecture and acetylcholinesterase-acetylcholine profile in Channa punctatus: amelioration by selenium. Comp Biochem Physiol C Toxicol Pharmacol. 144(1): 16–24. doi: 10.1016/j.cbpc.2006.04.018

Russ TC, Stamatakis E, Hamer M, et al. (2013) Socioeconomic status as a risk factor for dementia death: individual participant meta-analysis of 86 508 men and women from the UK. Br J Psychiatry. 203(1):10–7. doi: 10.1192/bjp.bp.112.119479.

Sabharwal V, Koushika SP. (2019) Crowd Control: Effects of Physical Crowding on Cargo Movement in Healthy and Diseased Neurons. Front Cell Neurosci. 13: 470. doi: 10.3389/fncel.2019.00470.

Sakaniwa R, Shirai K, Cadar D, et al. (2024) Socioeconomic Status Transition Throughout Life and Risk of Dementia. JAMA Netw Open. 7(5):e2412303. doi: 10.1001/jamanetworkopen.2024.12303

Saykin AJ, Wishart HA, Rabin LA, et al. (2006) Older adults with cognitive complaints show brain atrophy similar to that of amnestic MCI. Neurology. 67(5): 834–42. doi: 10.1212/01.wnl.0000234032.77541.a2.

Schuff N, Woerner N, Boreta L. et al. (2009) MRI of hippocampal volume loss in early Alzheimer’s disease in relation to ApoE genotype and biomarkers. Brain. 132(4): 1067–1077 doi: 10.1093/brain/awp007

Shah RC, Wilson RS, Tang Y, et al. (2009) Relation of hemoglobin to level of cognitive function in older persons. Neuroepidemiology. 32(1): 40–6. doi: 10.1159/000170905.

Smith S.M. (2002) Fast robust automated brain extraction. Human Brain Mapping. 17(3):143–155.

Spence H, McNeil CJ, Waiter GD. (2022) Cognition and brain iron deposition in whole grey matter regions and hippocampal subfields. Eur J Neurosci. 56(11):6039–6054. doi: 10.1111/ejn.15838

Spence H, Mengoa-Fleming S, Sneddon AA, et al. (2024) Associations between sex, systemic iron and inflammatory status and subcortical brain iron. Eur J Neurosci. 60(5):5069–5085. doi: 10.1111/ejn.16467.

Tachibana A, Iga J, Ozaki T, et al. (2024) Serum high-sensitivity C-reactive protein and dementia in a community-dwelling Japanese older population (JPSC-AD). Sci Rep 14:7374. doi: 10.1038/s41598-024-57922-1

Tacu I, Kokalari I, Abollino O, et al. (2021). Mechanistic Insights into the Role of Iron, Copper, and Carbonaceous Component on the Oxidative Potential of Ultrafine Particulate Matter. Chemical Research in Toxicology. 34(3): 767–779. doi: 10.1021/acs.chemrestox.0c00399

Tchounwou PB, Yedjou CG, Patlolla AK, Sutton DJ. (2012) Heavy metal toxicity and the environment. Exp Suppl. 101: 133–64. doi: 10.1007/978-3-7643-8340-4_6.

Tonne C, Elbaz A, Beevers S, Singh-Manoux A. (2014) Traffic-related air pollution in relation to cognitive function in older adults. Epidemiology 25:674–681. doi: 10.1097/EDE.0000000000000144

Tsagatakis I, Richardson J, Evangelides C, et al. (2023) UK Spatial Emissions Methodology: A report of the National Atmospheric Emission Inventory 2021. Retrieved from: https://naei.beis.gov.uk/reports/reports?report_id=1112 https://naei.energysecurity.gov.uk/reports/ uk-spatial-emissions-methodology-report-national-atmospheric-emission-inventory-2021

Van Der Maas HLJ, Kan KJ, Marsman M, Stevenson CE. (2017) Network Models for Cognitive Development and Intelligence. J Intell. 5(2):16. doi: 10.3390/jintelligence5020016.

Wei T, Simko V (2024). R package ‘corrplot’: Visualization of a Correlation Matrix. (Version 0.95), https://github.com/taiyun/corrplot.

Wickham H (2016). ggplot2: Elegant Graphics for Data Analysis. Springer-Verlag New York. ISBN 978-3-319-24277-4, https://ggplot2.tidyverse.org.

Wickham, H., Vaughan, D., & Girlich, M. (2024). tidyr: Tidy Messy Data. R package version 1.3.1. https://github.com/tidyverse/tidyr, https://tidyr.tidyverse.org.

Wolters FJ, Zonneveld HI, Licher S, et al. (2019) Hemoglobin and anemia in relation to dementia risk and accompanying changes on brain MRI. Neurology. 93: e9176. doi: 10.1212/WNL.0000000000008003

Xue M, Xu W, Ou YN, et al. (2019) Diabetes mellitus and risks of cognitive impairment and dementia: A systematic review and meta-analysis of 144 prospective studies. Ageing Res Rev.55: 100944. doi: 10.1016/j.arr.2019.100944.

Zhang Y, Chen SD, Deng YT, et al. (2023) Identifying modifiable factors and their joint effect on dementia risk in the UK Biobank. Nat Hum Behav. 7(7):1185–1195. doi: 10.1038/s41562-023-01585-x.

