## Supplementary Material for "Machine learning identifies routine blood tests as accurate predictive measures of pollution-dependent poor cognitive function"

**Supplementary Table 1.** Annual limits for ambient atmospheric pollutant measures. [a] Total days of exposure where mean concentrations are ≥120 µg $m^{-3}$

| | Variable | Metric | Units | Start | End | Annual limit |
|---|---|---|---|---|---|---|
| DEFRA modelled background atmospheric pollution data | Benzene | Annual mean | µg $m^{-3}$ | 2003 | 2022 | 5 µg $m^{-3}$ |
| | Carbon monoxide | Annual mean | mg $m^{-3}$ | 2002 | 2010 | 10 mg $m^{-3}$ |
| | | Maximum 8-hour mean | mg $m^{-3}$ | 2002 | 2010 | - |
| | $NO_2$ | Annual mean | µg $m^{-3}$ | 2001 | 2022 | 40 µg $m^{-3}$ |
| | $NO_X$ | Annual mean | µg $m^{-3}$ | 2001 | 2022 | 40 µg $m^{-3}$ |
| | Ozone | Total[a] | Days | 2003 | 2022 | 25 days |
| | $PM_{2.5}$ | Annual mean | µg $m^{-3}$ | 2002 | 2022 | 25 µg $m^{-3}$ |
| | $SO_2$ | Annual mean | µg $m^{-3}$ | 2002 | 2022 | 125 µg $m^{-3}$ |
| NAEI total atmospheric pollutants emissions | Arsenic | Total | kg/$km^2$ | 2010 | 2020 | 6 µg $m^{-3}$ |
| | Cadmium | Total | kg/$km^2$ | 2010 | 2020 | 5 µg $m^{-3}$ |
| | Chromium | Total | kg/$km^2$ | 2010 | 2020 | - |
| | Copper | Total | kg/$km^2$ | 2010 | 2020 | - |
| | Lead | Total | kg/$km^2$ | 2010 | 2020 | 0.5 µg $m^{-3}$ |
| | Mercury | Total | kg/$km^2$ | 2010 | 2020 | - |
| | Nickel | Total | kg/$km^2$ | 2010 | 2020 | 20 µg $m^{-3}$ |
| | Selenium | Total | kg/$km^2$ | 2010 | 2020 | - |
| | Vanadium | Total | kg/$km^2$ | 2010 | 2020 | - |
| | Zinc | Total | kg/$km^2$ | 2010 | 2020 | - |

**Supplementary Table 2.** All features that were included in the random forest model. Purple represents socioeconomic and demographic factors, green represents pollution exposure measures, Blue represents imaging measures and red represents blood markers.

| Feature |
|---|
| Education |
| Infection |
| Age |
| Sex |
| Height |
| Weight |
| BMI |
| Total Population Estimate |
| Working Age Population |
| Overal SIMD16 Rank |
| SIMD 2016 Percentile |
| SIMD 2016 Vigintile |
| SIMD 2016 Decile |
| SIMD 2016 Quintile |
| Income Domain 2016 Rank |
| Employment Domain 2016 Rank (SIMD) |
| Health Domain 2016 Rank (SIMD) |
| Education Domain 2016 Rank (SIMD) |
| Geographic Access Domain 2016 Rank (SIMD) |
| Crime Domain 2016 Rank (SIMD) |
| Housing Domain 2016 Rank (SIMD) |
| Income Rate (Percentage of income deprived people) |
| Income Count (Number of income deprived people) |
| Employment Rate (Percentage of employment deprived people) |
| Employment Count (Number of employment deprived people) |
| CIF (Comparative Illness Factor: standardised ratio) |
| ALCOHOL (Hospital stays related to alcohol use: standardised ratio) |
| DRUG (Hospital stays related to drug use: standardised ratio ) |
| SMR (Standardised mortality ratio) |
| DEPRESS (Proportion of population prescribed drugs for anxiety, depression or psychosis) |
| LBWT (Proportion of live singleton births of low birth weight) |
| EMERG (Emergency stays in hospital: standardised ratio) |
| School Attendance |
| School Attainment |
| Noquals (Working age people with no qualifications: standardised ratio ) |
| NEET (young people not in education, employment or training) |
| HESA (Higher Education Statistics Agency - SIMD) |
| Driving Travel Time To Petrol |
| Driving Travel Time To GP |
| Driving Travel Time To Post Office |
| Driving Travel Time To Primary School |
| Driving Travel Time To Retail |
| Driving Travel Time To Secondary School |
| Public Transport Travel Time to GP |
| Public Transport Travel Time to Post Office |
| Public Transport Travel Time to Retail |
| Crime Rate |
| Number of People in Overcrowded Households |
| Number of Houses with Non-central Heating |

| |
|---|
| Percentage of People in Overcrowded Households |
| Percentage of Houses with Non-central Heating |
| SO2_2002 |
| SO2_2003 |
| SO2_2004 |
| SO2_2005 |
| SO2_2006 |
| SO2_2007 |
| SO2_2008 |
| SO2_2009 |
| SO2_2010 |
| SO2_2011 |
| SO2_2012 |
| SO2_2013 |
| SO2_2014 |
| SO2_2015 |
| SO2_2016 |
| SO2_2017 |
| SO2_2018 |
| SO2_2019 |
| SO2_2020 |
| SO2_2021 |
| SO2_2022 |
| Benzene_2003 |
| Benzene_2004 |
| Benzene_2005 |
| Benzene_2006 |
| Benzene_2007 |
| Benzene_2008 |
| Benzene_2009 |
| Benzene_2010 |
| Benzene_2011 |
| Benzene_2012 |
| Benzene_2013 |
| Benzene_2014 |
| Benzene_2015 |
| Benzene_2016 |
| Benzene_2017 |
| Benzene_2018 |
| Benzene_2019 |
| Benzene_2020 |
| Benzene_2021 |
| Benzene_2022 |
| NO2_2001 |
| NO2_2002 |
| NO2_2003 |
| NO2_2004 |
| NO2_2005 |
| NO2_2006 |
| NO2_2007 |

| |
|---|
| NO2_2008 |
| NO2_2009 |
| NO2_2010 |
| NO2_2011 |
| NO2_2012 |
| NO2_2013 |
| NO2_2014 |
| NO2_2015 |
| NO2_2016 |
| NO2_2017 |
| NO2_2018 |
| NO2_2019 |
| NO2_2020 |
| NO2_2021 |
| NO2_2022 |
| NOX_2001 |
| NOX_2002 |
| NOX_2003 |
| NOX_2004 |
| NOX_2005 |
| NOX_2006 |
| NOX_2007 |
| NOX_2008 |
| NOX_2009 |
| NOX_2010 |
| NOX_2011 |
| NOX_2012 |
| NOX_2013 |
| NOX_2014 |
| NOX_2015 |
| NOX_2016 |
| NOX_2017 |
| NOX_2018 |
| NOX_2019 |
| NOX_2020 |
| NOX_2021 |
| NOX_2022 |
| OZONE_2003 |
| OZONE_2004 |
| OZONE_2005 |
| OZONE_2006 |
| OZONE_2007 |
| OZONE_2008 |
| OZONE_2009 |
| OZONE_2010 |
| OZONE_2011 |
| OZONE_2012 |
| OZONE_2013 |
| OZONE_2014 |
| OZONE_2015 |

| |
|---|
| OZONE_2016 |
| OZONE_2017 |
| OZONE_2018 |
| OZONE_2019 |
| OZONE_2020 |
| OZONE_2021 |
| OZONE_2022 |
| PM25_2002 |
| PM25_2003 |
| PM25_2004 |
| PM25_2005 |
| PM25_2006 |
| PM25_2007 |
| PM25_2008 |
| PM25_2009 |
| PM25_2010 |
| PM25_2011 |
| PM25_2012 |
| PM25_2013 |
| PM25_2014 |
| PM25_2015 |
| PM25_2016 |
| PM25_2017 |
| PM25_2018 |
| PM25_2019 |
| PM25_2020 |
| PM25_2021 |
| PM25_2022 |
| CO_2002 |
| CO_2003 |
| CO_2004 |
| CO_2005 |
| CO_2006 |
| CO_2007 |
| CO_2008 |
| CO_2009 |
| CO_2010 |
| CO (MAX) 2002 |
| CO (MAX) 2003 |
| CO (MAX) 2004 |
| CO (MAX) 2005 |
| CO (MAX) 2006 |
| CO (MAX) 2007 |
| CO (MAX) 2008 |
| CO (MAX) 2009 |
| CO (MAX) 2010 |
| Arsenic_2010 |
| Arsenic_2011 |
| Arsenic_2012 |
| Arsenic_2013 |

| |
|---|
| Arsenic_2014 |
| Arsenic_2015 |
| Arsenic_2016 |
| Arsenic_2017 |
| Arsenic_2018 |
| Arsenic_2019 |
| Arsenic_2020 |
| Cadmium_2010 |
| Cadmium_2011 |
| Cadmium_2012 |
| Cadmium_2013 |
| Cadmium_2014 |
| Cadmium_2015 |
| Cadmium_2016 |
| Cadmium_2017 |
| Cadmium_2018 |
| Cadmium_2019 |
| Cadmium_2020 |
| Chromium_2010 |
| Chromium_2011 |
| Chromium_2012 |
| Chromium_2013 |
| Chromium_2014 |
| Chromium_2015 |
| Chromium_2016 |
| Chromium_2017 |
| Chromium_2018 |
| Chromium_2019 |
| Chromium_2020 |
| Copper_2010 |
| Copper_2011 |
| Copper_2012 |
| Copper_2013 |
| Copper_2014 |
| Copper_2015 |
| Copper_2016 |
| Copper_2017 |
| Copper_2018 |
| Copper_2019 |
| Copper_2020 |
| Lead_2010 |
| Lead_2011 |
| Lead_2012 |
| Lead_2013 |
| Lead_2014 |
| Lead_2015 |
| Lead_2016 |
| Lead_2017 |
| Lead_2018 |
| Lead_2019 |

| |
|---|
| Lead_2020 |
| Mercury_2010 |
| Mercury_2011 |
| Mercury_2012 |
| Mercury_2013 |
| Mercury_2014 |
| Mercury_2015 |
| Mercury_2016 |
| Mercury_2017 |
| Mercury_2018 |
| Mercury_2019 |
| Mercury_2020 |
| Nickel_2010 |
| Nickel_2011 |
| Nickel_2012 |
| Nickel_2013 |
| Nickel_2014 |
| Nickel_2015 |
| Nickel_2016 |
| Nickel_2017 |
| Nickel_2018 |
| Nickel_2019 |
| Nickel_2020 |
| Selenium_2010 |
| Selenium_2011 |
| Selenium_2012 |
| Selenium_2013 |
| Selenium_2014 |
| Selenium_2015 |
| Selenium_2016 |
| Selenium_2017 |
| Selenium_2018 |
| Selenium_2019 |
| Selenium_2020 |
| Vanadium_2010 |
| Vanadium_2011 |
| Vanadium_2012 |
| Vanadium_2013 |
| Vanadium_2014 |
| Vanadium_2015 |
| Vanadium_2016 |
| Vanadium_2017 |
| Vanadium_2018 |
| Vanadium_2019 |
| Vanadium_2020 |
| Zinc_2010 |
| Zinc_2011 |
| Zinc_2012 |
| Zinc_2013 |
| Zinc_2014 |
| Zinc_2015 |

| |
|---|
| Zinc_2016 |
| Zinc_2017 |
| Zinc_2018 |
| Zinc_2019 |
| Zinc_2020 |
| Left Thalamus Iron |
| Left Caudate Iron |
| Left Putamen Iron |
| Left Pallidum Iron |
| Left Hippocampus Iron |
| Left Amygdala Iron |
| Right Thalamus Iron |
| Right Caudate Iron |
| Right Putamen Iron |
| Right Pallidum Iron |
| Right Hippocampus Iron |
| Right Amygdala Iron |
| Left Hippocampus Volume |
| Left Amygdala Volume |
| Left Thalamus Volume |
| Left Caudate Volume |
| Left Putamen Volume |
| Left Pallidum Volume |
| Right Thalamus Volume |
| Right Caudate Volume |
| Right Putamen Volume |
| Right Pallidum Volume |
| Right Hippocampus Volume |
| Right Amygdala Volume |
| Left Cerebral White Matter Volume |
| Left Lateral Ventricle Volume |
| Left Cerebellum White Matter Volume |
| Left Cerebellum Cortex Volume |
| 4th Ventricle Volume |
| Brain Stem Volume |
| Cerebral Spinal Fluid Volume |
| Left Accumbens Area Volume |
| Left Ventral Diencephalon Volume |
| Right Cerebral White Matter Volume |
| Right Lateral Ventricle Volume |
| Right Cerebellum White Matter Volume |
| Right Cerebellum Cortex Volume |
| Right Accumbens Area Volume |
| Right Ventral Diencephalon Volume |
| White Matter Hypointensities Volume |
| Corpus Callossum (Posterior) Volume |
| Corpus Callossum (Mid-Posterior) Volume |
| Corpus Callossum (Central) Volume |
| Corpus Callossum (Mid-Anterior) Volume |

| |
|---|
| Corpus Callossum (Anterior) Volume |
| Left Bankssts Cortex Volume |
| Left Caudal Anterior Cingulate Cortex Volume |
| Left Caudal middle frontal Cortex Volume |
| Left Cuneus Cortex Volume |
| Left Entorhinal Cortex Volume |
| Left Fusiform Cortex Volume |
| Left Inferior Parietal Cortex Volume |
| Left Inferior Temporal Cortex Volume |
| Left Isthmus Cingulate Cortex Volume |
| Left Lateral Occipital Cortex Volume |
| Left Lateral Orbitofrontal Cortex Volume |
| Left Lingual Cortex Volume |
| Left Medialo Orbitofrontal Cortex Volume |
| Left Middle Temporal Cortex Volume |
| Left Parahippocampal Cortex Volume |
| Left Paracentral Cortex Volume |
| Left Parsopercularis Volume |
| Left Parsorbitalis Volume |
| Left Parstriangularis Volume |
| Left Pericalcarine Cortex Volume |
| Left Postcentral Cortex Volume |
| Left Posterior Cingulate Cortex Volume |
| Left Precentral Cortex Volume |
| Left Precuneus Cortex Volume |
| Left Rostral Anterior Cingulate Cortex Volume |
| Left Rostral Middle Frontal Cortex Volume |
| Left Superior Frontal Cortex Volume |
| Left Superior Parietal Cortex Volume |
| Left Superior Temporal Cortex Volume |
| Left Supramarginal Cortex Volume |
| Left Frontalp Pole Volume |
| Left Temporal Pole Volume |
| Left Insula Cortex Volume |
| Right Bankssts Cortex Volume |
| Right Caudal Anterior Cingulate Cortex Volume |
| Right Caudal middle frontal Cortex Volume |
| Right Cuneus Cortex Volume |
| Right Entorhinal Cortex Volume |
| Right Fusiform Cortex Volume |
| Right Inferior Parietal Cortex Volume |
| Right Inferior Temporal Cortex Volume |
| Right Isthmus Cingulate Cortex Volume |
| Right Lateral Occipital Cortex Volume |
| Right Lateral Orbitofrontal Cortex Volume |
| Right Lingual Cortex Volume |
| Right Medialo Orbitofrontal Cortex Volume |
| Right Middle Temporal Cortex Volume |
| Right Parahippocampal Cortex Volume |
| Right Paracentral Cortex Volume |
| Right Parsopercularis Volume |

| |
|---|
| Right Parsorbitalis Volume |
| Right Parstriangularis Volume |
| Right Pericalcarine Cortex Volume |
| Right Postcentral Cortex Volume |
| Right Posterior Cingulate Cortex Volume |
| Right Precentral Cortex Volume |
| Right Precuneus Cortex Volume |
| Right Rostral Anterior Cingulate Cortex Volume |
| Right Rostral Middle Frontal Cortex Volume |
| Right Superior Frontal Cortex Volume |
| Right Superior Parietal Cortex Volume |
| Right Superior Temporal Cortex Volume |
| Right Supramarginal Cortex Volume |
| Right Frontalp Pole Volume |
| Right Temporal Pole Volume |
| Right Insula Cortex Volume |
| 3rd Ventricle Volume |
| Left Choroid Plexus Volume |
| Right Choroid Plexus Volume |
| Right Transverse Temporal Cortex Volume |
| Left Transverse Temporal Cortex Volume |
| Plasma Ferritin |
| Plasma Soluble Transferrin Receptor |
| Plasma Macrophage Colony Stimulating Factor |
| Plasma IL6 |
| Plasma IL1B |
| Grip Strength |
| C-reactive Protein |
| Haemoglobin |
| Red Blood Cell Count |
| Haematocrit |
| Mean Corpuscular Volume |
| Mean Corpuscular Haemoglobin |
| Platelet Count |
| White Blood Cell Count |
| Neutrophil Count |
| Eosinophil Count |
| Basophil Count |
| Lymphocyte Count |
| Monocyte Count |
| Large Unstained Cell Count |
| Mean Corpuscular Haemoglobin Concentration |

**Supplementary Table 3.** Random forest leave-one-out feature analysis for g score showing the proportion correct when a feature was left out of the model. Features that worsened predictivity of the model when left out were considered predictors of cognitive performance. Purple represents socioeconomic and demographic factors, green represents pollution exposure measures, Blue represents imaging measures and red represents blood markers.

| Feature | % Improvement |
|---|---|
| Age | 3.08 |
| School Attainment | 3.08 |
| Sex | 3.08 |
| Number of Housese with Non-central Heating | 3.08 |
| Arsenic (2012) | 3.08 |
| Arsenic (2013) | 3.08 |
| Arsenic (2014) | 3.08 |
| Arsenic (2015) | 3.08 |
| Arsenic (2020) | 3.08 |
| Carbon Monoxide (MAX; 2006) | 3.08 |
| Carbon Monoxide (MAX; 2007) | 3.08 |
| Carbon Monoxide (MAX; 2008) | 3.08 |
| Cadmium (2011) | 3.08 |
| Cadmium (2012) | 3.08 |
| Cadmium (2014) | 3.08 |
| Cadmium (2016) | 3.08 |
| Copper (2014) | 3.08 |
| Copper (2016) | 3.08 |
| Copper (2017) | 3.08 |
| Copper (2018) | 3.08 |
| Copper (2019) | 3.08 |
| Lead (2010) | 3.08 |
| Lead (2011) | 3.08 |
| Lead (2013) | 3.08 |
| Mercury (2011) | 3.08 |
| Mercury (2012) | 3.08 |
| Mercury (2017) | 3.08 |
| Mercury (2018) | 3.08 |
| Particulate Matter (PM2.5; 2005) | 3.08 |
| Particulate Matter (PM2.5; 2009) | 3.08 |
| Particulate Matter (PM2.5; 2017) | 3.08 |
| Particulate Matter (PM2.5; 2019) | 3.08 |
| Particulate Matter (PM2.5; 2022) | 3.08 |
| Sulfur Dioxide (2003) | 3.08 |
| Sulfur Dioxide (2010) | 3.08 |
| Corpus Callossum Central volume | 3.08 |
| Corpus Callossum Mid-Posterior volume | 3.08 |

| | |
|---|---|
| Left Pallidum Iron | 3.08 |
| Left Putamen Iron | 3.08 |
| Left Accumbens Area Volume | 3.08 |
| Left Amygdala Volume | 3.08 |
| Left Hippocampus Volume | 3.08 |
| Left Thalamus Volume | 3.08 |
| 4th Ventricle Volume | 3.08 |
| Left Caudalanteriorcingulate Cortex Volume | 3.08 |
| Left Cuneus Volume | 3.08 |
| Left Inferiortemporal Cortex Volume | 3.08 |
| Left Lingual Cortex Volume | 3.08 |
| Left Paracentral Cortex Volume | 3.08 |
| Left Parsopercularis Volume | 3.08 |
| Right Entorhinal Cortex Volume | 3.08 |
| C-Reactive Protein | 3.08 |
| White Blood Cell Count | 3.08 |
| Education | 6.08 |
| Weight (kg) | 6.08 |
| Rate of Overcrowding | 6.08 |
| Carbon Monoxide (2002) | 6.08 |
| Carbon Monoxide (2003) | 6.08 |
| Carbon Monoxide (2004) | 6.08 |
| Carbon Monoxide (2005) | 6.08 |
| Carbon Monoxide (2006) | 6.08 |
| Carbon Monoxide (2007) | 6.08 |
| Carbon Monoxide (MAX; 2009) | 6.08 |
| Carbon Monoxide (MAX; 2010) | 6.08 |
| Cadmium (2013) | 6.08 |
| Copper (2015) | 6.08 |
| Lead (2012) | 6.08 |
| Lead (2012) | 6.08 |
| Lead (2015) | 6.08 |
| Lead (2017) | 6.08 |
| Lead (2018) | 6.08 |
| Lead (2019) | 6.08 |
| Lead (2020) | 6.08 |
| Particulate Matter (PM2.5; 2007) | 6.08 |
| Particulate Matter (PM2.5; 2011) | 6.08 |
| Particulate Matter (PM2.5; 2018) | 6.08 |
| Corpus Callossum Anterior Volume | 6.08 |
| Corpus Callossum Mid-Anterior Volume | 6.08 |
| Corpus Callossum Posterior Volume | 6.08 |
| Left Ventral Diencephalon Volume | 6.08 |
| Right Amygdala Iron | 6.08 |
| Right Accumbens Area Volume | 6.08 |
| Right Cerebellum Cortex Volume | 6.08 |
| Right Cerebellum White Matter Volume | 6.08 |
| Right Pallidum Volume | 6.08 |

| | |
|---|---|
| Left Bankssts Cortex Volume | 6.08 |
| Left Caudal Middle Frontal Cortex Volume | 6.08 |
| Pericalcarine Cortex Volume | 6.08 |
| Left Postcentral Cortex Volume | 6.08 |
| Haematocrit | 6.08 |
| Haemoglobin | 6.08 |
| Mean Corpuscular Haemoglobin | 6.08 |
| Arsenic (2010) | 9.08 |
| Arsenic (2011) | 9.08 |
| Carbon Monoxide (2008) | 9.08 |
| Carbon Monoxide (2009) | 9.08 |
| Carbon Monoxide (2010) | 9.08 |
| Carbon Monoxide (MAX; 2002) | 9.08 |
| Carbon Monoxide (MAX; 2003) | 9.08 |
| Carbon Monoxide (MAX; 2004) | 9.08 |
| Carbon Monoxide (MAX; 2005) | 9.08 |
| Lead (2016) | 9.08 |
| Particulate Matter (PM2.5; 2010) | 9.08 |
| Particulate Matter (PM2.5; 2012) | 9.08 |
| Mean Corpuscular Volume | 12.08 |

**Supplementary Table 4.** Random forest leave-one-out feature analysis for memory score (immediate recall) showing the proportion correct when a feature was left out of the model. Features that worsened predictivity of the model when left out were considered predictors of cognitive performance. Purple represents socioeconomic and demographic factors, green represents pollution exposure measures, Blue represents imaging measures and red represents blood markers.

| Feature | % Improvement |
|---|---|
| Crime Domain 2016 Rank (SIMD) | 1.96 |
| EMERG (Emergency stays in hospital: standardised ratio) | 1.96 |
| Education Domain 2016 Rank (SIMD) | 1.96 |
| Geographic Domain 2016 Rank (SIMD) | 1.96 |
| Health Domain 2016 Rank (SIMD) | 1.96 |
| Income Domain 2016 Rank (SIMD) | 1.96 |
| LBWT (Proportion of live singleton births of low birth weight) | 1.96 |
| Overal SIMD16 Rank | 1.96 |
| SIMD 2016 Vigintile | 1.96 |
| SMR (Standardised mortality ratio) | 1.96 |
| Driving Travel Time To GP | 1.96 |
| Driving Travel Time To Post Office | 1.96 |
| Driving Travel Time To Primary School | 1.96 |
| Driving Travel Time To Retail | 1.96 |
| Number of People in Overcrowded Households | 1.96 |
| Arsenic (2011) | 1.96 |
| Arsenic (2012) | 1.96 |
| Arsenic (2016) | 1.96 |
| Arsenic (2017) | 1.96 |
| Arsenic (2018) | 1.96 |
| Arsenic (2019) | 1.96 |
| Arsenic (2020) | 1.96 |
| Benzene (2009) | 1.96 |
| Benzene (2014) | 1.96 |
| Benzene (2015) | 1.96 |
| Benzene (2016) | 1.96 |
| Benzene (2018) | 1.96 |
| Benzene (2020) | 1.96 |
| Benzene (2021) | 1.96 |
| Cadmium (2014) | 1.96 |
| Cadmium (2016) | 1.96 |
| Copper (2011) | 1.96 |
| Copper (2012) | 1.96 |
| Copper (2013) | 1.96 |
| Copper (2014) | 1.96 |
| NO2 (2017) | 1.96 |
| NOX (2002) | 1.96 |
| NOX (2003) | 1.96 |
| NOX (2005) | 1.96 |
| NOX (2009) | 1.96 |
| NOX (2010) | 1.96 |
| NOX (2011) | 1.96 |
| NOX (2012) | 1.96 |
| NOX (2013) | 1.96 |

| | |
|---|---|
| Nickel (2012) | 1.96 |
| Nickel (2013) | 1.96 |
| Nickel (2014) | 1.96 |
| Nickel (2016) | 1.96 |
| Nickel (2017) | 1.96 |
| Nickel (2018) | 1.96 |
| Nickel (2019) | 1.96 |
| Nickel (2020) | 1.96 |
| OZONE (2008) | 1.96 |
| OZONE (2016) | 1.96 |
| PM25 (2005) | 1.96 |
| PM25 (2014) | 1.96 |
| PM25 (2015) | 1.96 |
| PM25 (2017) | 1.96 |
| SO2 (2016) | 1.96 |
| Selenium (2010) | 1.96 |
| Selenium (2015) | 1.96 |
| Selenium (2016) | 1.96 |
| Selenium (2017) | 1.96 |
| Selenium (2018) | 1.96 |
| Selenium (2020) | 1.96 |
| Vanadium (2010) | 1.96 |
| Vanadium (2011) | 1.96 |
| Vanadium (2012) | 1.96 |
| Vanadium (2013) | 1.96 |
| Vanadium (2015) | 1.96 |
| Vanadium (2016) | 1.96 |
| Vanadium (2017) | 1.96 |
| Vanadium (2018) | 1.96 |
| Vanadium (2019) | 1.96 |
| Vanadium (2020) | 1.96 |
| Zinc (2010) | 1.96 |
| Zinc (2014) | 1.96 |
| Zinc (2015) | 1.96 |
| Zinc (2017) | 1.96 |
| Corpus Callossum (Central) Volume | 1.96 |
| Corpus Callossum (Mid-anterior) Volume | 1.96 |
| Corpus Callossum (Mid-posterior) Volume | 1.96 |
| Corpus Callossum (Posterior) Volume | 1.96 |
| Left Caudate Iron | 1.96 |
| Left Putamen Iron | 1.96 |
| Left Thalamus Iron | 1.96 |
| Left Caudate Volume | 1.96 |
| Left Hippocampus Volume | 1.96 |
| Right Amygdala Iron | 1.96 |
| Right Pallidum Iron | 1.96 |
| Right Cerebellum Cortex Volume | 1.96 |
| White Matter Hypointensities Volume | 1.96 |
| Left Paracentral Cortex Volume | 1.96 |

| | |
|---|---|
| Left Parsorbitalis Volume | 1.96 |
| Right Insula Cortex Volume | 1.96 |
| Right Paracentral Cortex Volume | 1.96 |
| Right Pericalcarine volume | 1.96 |
| C-reactive Protein | 1.96 |
| Lymphocyte Count | 1.96 |
| Percentage of Houses with Non-central Heating | 3.92 |
| Percentage of People in Overcrowded Households | 3.92 |
| School Pupil Attendance | 3.92 |
| Number of Employment Deprived People | 3.92 |
| HESA (Higher Education Statistics Agency - SIMD) | 3.92 |
| Housing Domain 2016 Rank | 3.92 |
| Public Transport Travel Time To GP | 3.92 |
| SIMD 2016 Percentile | 3.92 |
| Crime Rate | 3.92 |
| Benzene (2004) | 3.92 |
| Benzene (2005) | 3.92 |
| Benzene (2006) | 3.92 |
| Benzene (2007) | 3.92 |
| Benzene (2008) | 3.92 |
| Benzene (2010) | 3.92 |
| Benzene (2011) | 3.92 |
| Benzene (2012) | 3.92 |
| Cadmium (2013) | 3.92 |
| Mercury (2017) | 3.92 |
| Mercury (2018) | 3.92 |
| Mercury (2019) | 3.92 |
| Mercury (2020) | 3.92 |
| NO2 (2001) | 3.92 |
| NO2 (2018) | 3.92 |
| NO2 (2019) | 3.92 |
| NO2 (2020) | 3.92 |
| NOX (2006) | 3.92 |
| NOX (2007) | 3.92 |
| NOX (2008) | 3.92 |
| NOX (2014) | 3.92 |
| NOX (2015) | 3.92 |
| Nickel (2011) | 3.92 |
| OZONE (2007) | 3.92 |
| OZONE (2011) | 3.92 |
| OZONE (2015) | 3.92 |
| SO2 (2017) | 3.92 |
| SO2 (2018) | 3.92 |
| SO2 (2022) | 3.92 |
| Vanadium (2014) | 3.92 |
| Zinc (2011) | 3.92 |
| Zinc (2012) | 3.92 |
| Zinc (2013) | 3.92 |

| | |
|---|---|
| Zinc (2016) | 3.92 |
| Zinc (2018) | 3.92 |
| Zinc (2019) | 3.92 |
| Zinc (2020) | 3.92 |
| Cerebral Spinal Fluid Volume | 3.92 |
| Left Accumbens Area Volume | 3.92 |
| Left Amygdala Volume | 3.92 |
| Left Cerebellum White Matter Volume | 3.92 |
| Left Inferior Lateral Ventricle Volume | 3.92 |
| Right Accumbens Area Volume | 3.92 |
| Right Cerebellum White Matter Volume | 3.92 |
| Right Cerebral White Matter Volume | 3.92 |
| Right Ventral Diencephalon Volume | 3.92 |
| Left Medial Orbitofrontal Cortex Volume | 3.92 |
| Left Parsopercularis Volume | 3.92 |
| Left Precuneus Cortex Volume | 3.92 |
| Left Rostral Anterior Cingulate Cortex Volume | 3.92 |
| Right Frontal Pole Volume | 3.92 |
| Right Lingual Cortex Volume | 3.92 |
| Right Parsorbitalis Volume | 3.92 |
| Right Postcentral Cortex Volume | 3.92 |
| Right Precentral Cortex Volume | 3.92 |
| Right Superior Frontal Cortex Volume | 3.92 |
| Right Superior Parietal Cortex Volume | 3.92 |
| Right Temporal Pole Volume | 3.92 |
| Plasma Lead (208 Pb [ No Gas ]) | 3.92 |
| Haemoglobin | 3.92 |
| Council Area (SIMD) | 5.88 |
| DEPRESS (Proportion of population prescribed drugs for anxiety, depression or psychosis) | 5.88 |
| Employment Rate (Percentage of employment deprived people) | 5.88 |
| Income Rate (Percentage of income deprived people) | 5.88 |
| Public Transport Travel Time To Retail | 5.88 |
| Total Population Estimation (SIMD) | 5.88 |
| Driving Travel Time To Secondary School | 5.88 |
| Number of People in Houses with Non-central Heating | 5.88 |
| Benzene (2003) | 5.88 |
| Cadmium (2011) | 5.88 |
| Mercury (2016) | 5.88 |
| NO2 (2006) | 5.88 |
| NO2 (2008) | 5.88 |
| NO2 (2009) | 5.88 |
| NO2 (2010) | 5.88 |
| NO2 (2011) | 5.88 |
| NO2 (2012) | 5.88 |
| NO2 (2013) | 5.88 |
| NO2 (2014) | 5.88 |
| NO2 (2015) | 5.88 |
| NO2 (2016) | 5.88 |

| | |
|---|---|
| NOX (2004) | 5.88 |
| NOX (2017) | 5.88 |
| Nickel (2010) | 5.88 |
| OZONE (2006) | 5.88 |
| SO2 (2019) | 5.88 |
| SO2 (2021) | 5.88 |
| Left Amygdala Iron | 5.88 |
| Left Putamen Volume | 5.88 |
| Left Ventral Diencephalon Volume | 5.88 |
| Right Thalamus Iron | 5.88 |
| Right Amygdala Volume | 5.88 |
| Right Caudate Volume | 5.88 |
| Right Lateral Ventricle Volume | 5.88 |
| Right Pallidum Volume | 5.88 |
| Right Putamen Volume | 5.88 |
| 4th Ventricle Volume | 5.88 |
| Left Parstriangularis Volume | 5.88 |
| Left Precentral Cortex Volume | 5.88 |
| Left Rostral Middle Frontal Cortex Volume | 5.88 |
| Left Superior Parietal Cortex Volume | 5.88 |
| Right Lateral Orbitofrontal Cortex Volume | 5.88 |
| Right Medial Orbitofrontal Cortex Volume | 5.88 |
| Right Middle Temporal Cortex Volume | 5.88 |
| Right Parahippocampal Cortex Volume | 5.88 |
| Right Parsopercularis Volume | 5.88 |
| Right Parstriangularis Volume | 5.88 |
| Right Posterior Cingulate Cortex Volume | 5.88 |
| Right Rostral Anterior Cingulate Cortex Volume | 5.88 |
| Right Rostral Middle Frontal Cortex Volume | 5.88 |
| Right Supramarginal Cortex Volume | 5.88 |
| Public Transport Travel Time To Post Office | 7.84 |
| DRUG (Hospital stays related to drug use: standardised ratio ) | 7.84 |
| NEET (young people not in education, employment or training) | 7.84 |
| Noquals (Working age people with no qualifications: standardised ratio ) | 7.84 |
| ALCOHOL (Hospital stays related to alcohol use: standardised ratio) | 7.84 |
| CIF (Comparative Illness Factor: standardised ratio) | 7.84 |
| Cadmium (2012) | 7.84 |
| NO2 (2007) | 7.84 |
| NOX (2016) | 7.84 |
| NOX (2018) | 7.84 |
| NOX (2019) | 7.84 |
| OZONE (2004) | 7.84 |
| SO2 (2020) | 7.84 |
| Left Hippocampus Iron | 7.84 |
| Left Pallidum Iron | 7.84 |
| Left Cerebral White Matter Volume | 7.84 |
| Left Lateral Ventricle Volume | 7.84 |
| Left Pallidum Volume | 7.84 |
| Right Thalamus Volume | 7.84 |

| | |
|---|---|
| Left Pericalcarine Cortex Volume | 7.84 |
| Left Posterior Cingulate Cortex Volume | 7.84 |
| Left Supramarginal Cortex Volume | 7.84 |
| Right Precuneus Cortex Volume | 7.84 |
| Right Superior Temporal Cortex Volume | 7.84 |
| Number of Income Deprived People | 9.8 |
| NOX (2020) | 9.8 |
| NOX (2021) | 9.8 |
| NOX (2022) | 9.8 |
| OZONE (2003) | 9.8 |
| Brain Stem Volume | 9.8 |
| Right Hippocampus Volume | 9.8 |
| Left Postcentral Cortex Volume | 9.8 |
| Left Superior Frontal Cortex Volume | 9.8 |
| Right Lateral Occipital Cortex Volume | 9.8 |
| Data Zone (SIMD) | 11.76 |
| Grip Strength | 11.76 |
| Intermediate Zone (SIMD) | 11.76 |
| Left Frontal Pole Volume | 11.76 |
| Right Caudal Middle Frontal Cortex Volume | 11.76 |
| Right Entorhinal Cortex Volume | 11.76 |
| Right Inferior Parietal Cortex Volume | 11.76 |
| Right Inferior Temporal Cortex Volume | 11.76 |
| Plasma Soluble Transferrin Receptor | 11.76 |
| Plasma IL6 | 11.76 |
| Plasma Macrophage Colony Stimulating Factor | 11.76 |
| Platelet Count | 11.76 |
| Plasma Calcium (44 Ca [ No Gas ]) | 11.76 |
| Plasma Copper (63 Cu [ He ]) | 11.76 |
| Plasma Iron (56 Fe [ He ]) | 11.76 |
| Plasma Magnesium (24 Mg [ He ]) | 11.76 |
| Plasma Phosphorus (31 P [ No Gas ]) | 11.76 |
| Plasma Potassium (39 K [ No Gas ]) | 11.76 |
| Plasma Selenium (78 Se [ H2 ]) | 11.76 |
| Plasma Zinc (66 Zn [ He ]) | 11.76 |
| Plasma Ferritin | 11.76 |
| Mean Corpuscular Haemoglobin Concentration | 11.76 |
| White Blood Cell Count | 11.76 |
| Left Superior Temporal Cortex Volume | 15.69 |
| Left Temporal Pole Volume | 15.69 |
| Right Fusiform Volume | 15.69 |
| Right Isthmus Cingulate Cortex Volume | 15.69 |
| Left Insula Cortex Volume | 17.65 |
| Right Bankssts Volume | 17.65 |
| Right Caudal Anterior Cingulate Cortex Volume | 17.65 |

**Supplementary Table 5.** Random forest leave-one-out feature analysis for memory score (delayed recall) showing the proportion correct when a feature was left out of the model. Features that worsened predictivity of the model when left out were considered predictors of cognitive performance. Purple represents socioeconomic and demographic factors, green represents pollution exposure measures, Blue represents imaging measures and red represents blood markers.

| Feature | % Improvement |
|---|---|
| Driving Travel Time To GP | 2.00 |
| Driving Travel Time To Secondary School | 2.00 |
| Income Domain 2016 Rank (SIMD) | 2.00 |
| Income Count (Number of income deprived people) | 2.00 |
| Overal SIMD16 Rank | 2.00 |
| SIMD 2016 Decile | 2.00 |
| SIMD 2016 Percentile | 2.00 |
| SIMD 2016 Vigintile | 2.00 |
| Arsenic (2012) | 2.00 |
| Arsenic (2016) | 2.00 |
| Arsenic (2017) | 2.00 |
| Arsenic (2018) | 2.00 |
| Arsenic (2019) | 2.00 |
| Arsenic (2020) | 2.00 |
| CO (2007) | 2.00 |
| CO MAX (2006) | 2.00 |
| Cadmium (2010) | 2.00 |
| Cadmium (2011) | 2.00 |
| Cadmium (2018) | 2.00 |
| Chromium (2010) | 2.00 |
| Lead (2015) | 2.00 |
| Lead (2016) | 2.00 |
| Lead (2017) | 2.00 |
| Lead (2018) | 2.00 |
| Lead (2019) | 2.00 |
| Mercury (2012) | 2.00 |
| Mercury (2016) | 2.00 |
| Mercury (2018) | 2.00 |
| Mercury (2020) | 2.00 |
| NO2 (2005) | 2.00 |
| NO2 (2008) | 2.00 |
| NO2 (2010) | 2.00 |
| NO2 (2011) | 2.00 |
| NO2 (2012) | 2.00 |
| NO2 (2013) | 2.00 |
| NOX (2021) | 2.00 |
| Nickel (2013) | 2.00 |
| Nickel (2019) | 2.00 |
| Nickel (2020) | 2.00 |
| OZONE (2007) | 2.00 |
| PM25 (2002) | 2.00 |
| PM25 (2003) | 2.00 |
| PM25 (2015) | 2.00 |

| | |
|---|---|
| SO2 (2006) | 2.00 |
| SO2 (2012) | 2.00 |
| Selenium (2011) | 2.00 |
| Selenium (2012) | 2.00 |
| Selenium (2013) | 2.00 |
| Selenium (2016) | 2.00 |
| Selenium (2019) | 2.00 |
| Selenium (2020) | 2.00 |
| Vanadium (2018) | 2.00 |
| Zinc (2012) | 2.00 |
| Zinc (2013) | 2.00 |
| Corpus Callossum (Posterior) Volume | 2.00 |
| Left Hippocampus Iron | 2.00 |
| Left Pallidum Iron | 2.00 |
| Left Amygdala Volume | 2.00 |
| Left Cerebellum Cortex Volume | 2.00 |
| Left Thalamus Volume | 2.00 |
| Right Amygdala Iron | 2.00 |
| Right Putamen Iron | 2.00 |
| Right Cerebellum Cortex Volume | 2.00 |
| Basophil Count | 2.00 |
| Driving Travel Time To Post Office | 4.00 |
| Driving Travel Time To Primary School | 4.00 |
| Housing Domain 2016 Rank (SIMD) | 4.00 |
| Income Rate (Percentage of income deprived people) | 4.00 |
| SIMD 2016 Quintile | 4.00 |
| Arsenic (2014) | 4.00 |
| Arsenic (2015) | 4.00 |
| CO (2006) | 4.00 |
| CO (2008) | 4.00 |
| Cadmium (2013) | 4.00 |
| Cadmium (2015) | 4.00 |
| Cadmium (2016) | 4.00 |
| Cadmium (2017) | 4.00 |
| Cadmium (2020) | 4.00 |
| Chromium (2011) | 4.00 |
| Mercury (2013) | 4.00 |
| Mercury (2015) | 4.00 |
| Mercury (2017) | 4.00 |
| Mercury (2019) | 4.00 |
| NO2 (2001) | 4.00 |
| NO2 (2002) | 4.00 |
| NO2 (2003) | 4.00 |
| NO2 (2004) | 4.00 |
| NO2 (2006) | 4.00 |
| Nickel (2010) | 4.00 |
| Nickel (2011) | 4.00 |
| Nickel (2012) | 4.00 |
| SO2 (2005) | 4.00 |

| | |
|---|---|
| Selenium (2010) | 4.00 |
| Selenium (2015) | 4.00 |
| Selenium (2018) | 4.00 |
| Vanadium (2014) | 4.00 |
| Vanadium (2015) | 4.00 |
| Vanadium (2016) | 4.00 |
| Vanadium (2017) | 4.00 |
| Vanadium (2019) | 4.00 |
| Vanadium (2020) | 4.00 |
| Zinc (2010) | 4.00 |
| Zinc (2011) | 4.00 |
| Zinc (2014) | 4.00 |
| Zinc (2015) | 4.00 |
| Zinc (2016) | 4.00 |
| Zinc (2017) | 4.00 |
| Zinc (2019) | 4.00 |
| Left Accumbens Area Volume | 4.00 |
| Left Pallidum Volume | 4.00 |
| Right Caudate Iron | 4.00 |
| C-Reactive Protein | 4.00 |
| Crime Domain 2016 Rank (SIMD) | 6.00 |
| Education Domain 2016 Rank (SIMD) | 6.00 |
| Employment Domain 2016 Rank (SIMD) | 6.00 |
| Employment Count (Number of employment deprived people) | 6.00 |
| Geographic Access Domain 2016 Rank (SIMD) | 6.00 |
| Health Domain 2016 Rank (SIMD) | 6.00 |
| Arsenic (2013) | 6.00 |
| CO (2010) | 6.00 |
| CO MAX (2007) | 6.00 |
| Cadmium (2014) | 6.00 |
| Cadmium (2019) | 6.00 |
| Chromium (2014) | 6.00 |
| Chromium (2016) | 6.00 |
| Chromium (2019) | 6.00 |
| Lead (2010) | 6.00 |
| Lead (2011) | 6.00 |
| Lead (2012) | 6.00 |
| Lead (2013) | 6.00 |
| NO2 (2007) | 6.00 |
| Zinc (2018) | 6.00 |
| Zinc (2020) | 6.00 |
| Left Ventral Diencephalon Volume | 6.00 |
| Right Cerebral White Matter Volume | 6.00 |
| Mean Corpuscular Haemoglobin | 6.00 |
| Total Population Estimation (SIMD) | 8.00 |
| Working Age Population | 8.00 |
| Driving Travel Time To Retail | 8.00 |
| Council Area (SIMD) | 8.00 |
| Employment Rate (Percentage of employment deprived people) | 8.00 |

| | |
|---|---|
| Benzene (2021) | 8.00 |
| Benzene (2022) | 8.00 |
| Chromium (2015) | 8.00 |
| Chromium (2017) | 8.00 |
| Chromium (2018) | 8.00 |
| Chromium (2020) | 8.00 |
| Copper (2011) | 8.00 |
| Copper (2012) | 8.00 |
| Copper (2019) | 8.00 |
| Selenium (2014) | 8.00 |
| Left Amygdala Iron | 8.00 |
| Right Thalamus Iron | 8.00 |
| Chromium (2012) | 10.00 |
| Chromium (2013) | 10.00 |
| Copper (2010) | 10.00 |
| Copper (2013) | 10.00 |
| Copper (2014) | 10.00 |
| Copper (2015) | 10.00 |
| Copper (2018) | 10.00 |
| Copper (2020) | 10.00 |
| Copper (2016) | 12.00 |
| Copper (2017) | 14.00 |

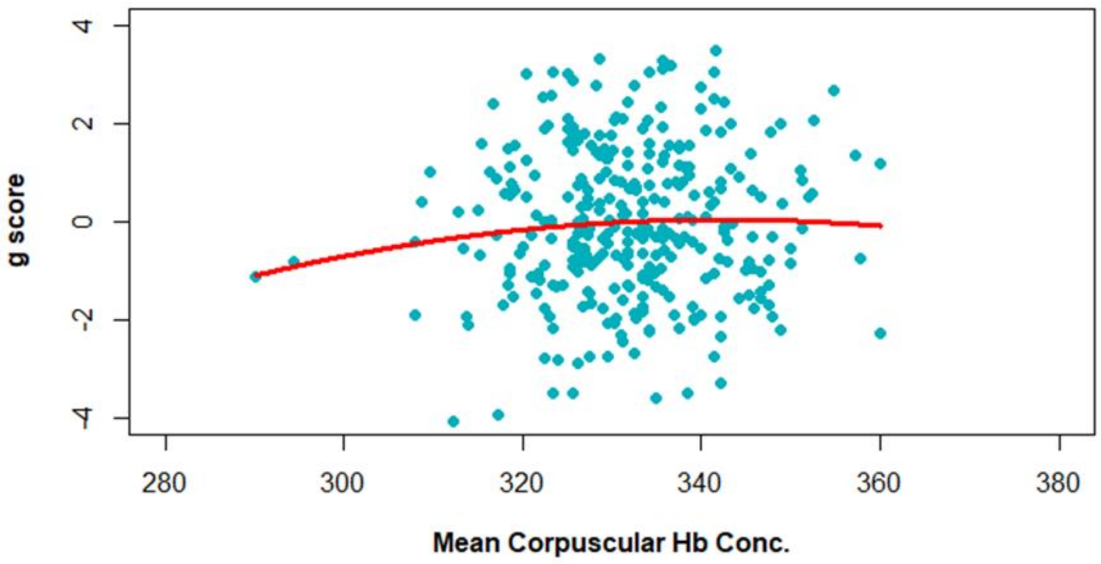

**Supplementary Figure 1.** Mean corpuscular haemoglobin concentration was not significantly associated with general cognition score (p=0.303).
